# Breaking boundaries: fungi in the “rhizoctonia” species complex exhibit systemic colonization in three terrestrial orchid species

**DOI:** 10.1101/2025.01.08.631981

**Authors:** Calevo Jacopo, Alibrandi Pasquale, Voyron Samuele, Girlanda Mariangela, Perotto Silvia

## Abstract

- Most green orchids associate with orchid mycorrhizal (OrM) fungi belonging to the ‘rhizoctonia’ complex, a polyphyletic group of Tulasnellaceae, Ceratobasidiaceae and Serendipitaceae (Agaricomycotina), which are generally assumed to live as saprotrophs in soil. However, OrM rhizoctonias were rarely detected by metabarcoding in soil around orchid roots, and we have tested the hypothesis that these fungi may use adult orchid plants as a niche by colonizing not only their roots, but also other organs.
- The occurrence of OrM rhizoctonias inside roots, stems and leaves of three terrestrial orchid species (*Spiranthes spiralis*, *Serapias vomeracea* and *Neottia ovata*) was therefore investigated by metabarcoding. To test the possibility of a vertical transmission of OrM fungi, reproductive structures (capsules, as well as seeds in *S. spiralis*) were also analyzed in a subset of plants.
- In all orchid species, a broad majority of OrM fungi found in roots was also detected in either stems or leaves of the same plant. OrM fungi were also detected in capsules/seeds.
- Systemic colonization of orchid tissues by OrM symbionts is a novel finding that raises important questions on the plant-fungus relationship in the aerial organs and opens intriguing perspectives on the potential modes of fungal transmission to the orchid progeny.

## Introduction

All plants share their lives with a complex and diverse microbial community that contributes to plant growth and defense against biotic and abiotic stress (Hardoim *et al*., 2015; Almario *et al*., 2017). Fungi are important components of the plant microbiota and can colonize the external surfaces as well as the internal tissues of all plant organs, both below-ground and above-ground (Choi *et al*., 2021). The root-associated mycobiota comprises mycorrhizal fungi, a diverse group of symbiotic fungi that form specialized plant-fungus interfaces inside the roots of most land plants (Smith & Read, 2008). Mycorrhizal symbioses are thought to have been instrumental for land colonization by early plants, and their ecological success and wide distribution are likely related to the improved mineral nutrition and health of the host plant (Genre *et al*., 2020). Most plants benefit from the mycorrhizal association, some plant species, like orchids, being particularly reliant on these symbiotic fungi for their survival.

Orchids belong to one of the largest families of flowering plants, counting more than 28,000 terrestrial and epiphytic species distributed in extremely diverse habitats, ranging from tropical forests to semi-arid deserts (Chase *et al*., 2015; Christenhusz & Byng, 2016). Despite their diversity, survival of all orchids in nature is intrinsically bound to the association with orchid mycorrhizal (OrM) fungi. All orchids produce tiny “dust” seeds devoid of stored nutrients and with an immature embryo (Arditti & Ghani, 2000). Orchid seeds that successfully germinate develop a protocorm, a postembryonic heterotrophic structure that precedes seedling formation (Arditti & Ghani, 2000; Rasmussen, 1995). OrM fungi are necessary in these early stages because they induce seed germination and provide germinating seeds and protocorms with organic carbon and other essential nutrients (Smith & Read, 2008; De Rose et al., 2023). Mycorrhizal orchid protocorms eventually develop into adult plants by forming aerial organs that can be achlorophyllous or chlorophyllous. Achlorophyllous orchids lack photosynthesis and remain completely dependent on their OrM fungal symbionts for organic carbon supply, a strategy termed full mycoheterotrophy (Hynson *et al*., 2013). However, stable isotope natural abundance indicates that green orchids can also receive carbon from their symbiotic fungal partners. In particular, green orchids living in shady forest habitats can supplement inefficient photosynthesis with fungal-derived carbon, a strategy termed partial mycoheterotrophy or mixotrophy (Selosse & Roy, 2009; Merckx, 2013). Even photosynthetic orchids living in open meadows, expected to be fully autotrophic, can use OrM fungi to supplement their carbon demands in some cases (Gebauer *et al*., 2016; Girlanda *et al*., 2011; Schiebold *et al*., 2018). Thus, OrM fungi are critically important for seed germination, seedling survival and plant growth in nature, and it is therefore not surprising that local-scale distribution and population dynamics of orchids can be limited by distribution and abundance of their symbiotic OrM fungi (McCormick & Jacquemyn, 2014; McCormick *et al*., 2018).

OrM fungi belong to diverse taxa, whose phylogenetic position mirrors the habitat and the trophic abilities of the host plants. Terrestrial achlorophyllous and green orchids restricted to forest floors usually form specific associations with fungi also capable of forming ectomycorrhiza (ECM) on photosynthetic trees (Dearnaley *et al*., 2012), but associations with wood-decomposers have been also reported (Ogura-Tsujita *et al*., 2021). Photoautotrophic orchids mainly associate with OrM fungi belonging to the ‘rhizoctonia’ species complex, a polyphyletic group comprising teleomorphs in three distinct families of Agaricomycotina (Basidiomycota): Tulasnellaceae and Ceratobasidiaceae in the Cantharellales, and Serendipitaceae in the Sebacinales (Roberts, 1999; Taylor *et al*., 2002; Weiß *et al*., 2004, 2016). The biology and ecology of rhizoctonia-like OrM fungi are poorly understood when compared with ECM fungi, also because of their inconspicuous nature (Roberts, 1999). An intriguing feature of rhizoctonia-like OrM fungi is that some members are not uniquely mycorrhizal, as they have been found as non-mycorrhizal endophytes in the roots of non-orchid plants, where they can promote plant growth and tolerance to biotic and abiotic stress (Lahrmann *et al*., 2015; Sarkar *et al*., 2019; Mahdi *et al*., 2022; Mosquera-Espinosa *et al*., 2013; Ray *et al*., 2018). For example, the OrM fungus *Serendipita* (= *Sebacina* or *Piriformospora*) *vermifera*, originally isolated from mycorrhizal roots of terrestrial Australian orchids in the genus *Caladenia* (Warcup and & Talbot, 1967), is considered a generalist root endophyte because it can colonize a wide range of monocots and dicots without forming typical mycorrhizal fungal coils (Weiß *et al*., 2016; Qiang *et al*., 2012; Unnikumar *et al*., 2013). A related species, *Serendipita indica*, improves the availability of nitrogen (Saleem *et al*., 2022) and produces phosphatases and organic acids which contribute to solubilization of phosphate from insoluble polyphosphates and organic phosphates (Johnson *et al*., 2014). In soybean, *S. indica* improved N, P and K uptake as a result of upregulation of transporter genes related to P and N uptake(Bajaj *et al*., 2018). It also produces plant growth regulators that affect the root architecture, such as indole acetic acid (IAA, the most common natural auxin) and cytokinins, which have a positive effect on root growth (Oelmüller *et al*., 2009; Liu *et al*., 2023). Other OrM rhizoctonia-like fungi have been found in the roots of non-orchid plants, although their potential role in non-orchid hosts is unknown (Selosse *et al*., 2022). For example, the same *Tulasnella* sp. OTU (Operational Taxonomic Unit) was identified in the roots of *Orchis purpurea* and of a nearby *Bromus erectus* plant in a Mediterranean meadow (Girlanda *et al*., 2011).

Saprotrophic capabilities have been often reported for OrM rhizoctonia-like fungi grown *in vitro*, where these fungi can produce a range of plant cell wall degrading enzymes (Nurfadilah *et al*., 2013, Zhao *et al*., 2021; Novotná *et al*., 2023). In agreement with these observations, the sequenced genome of two OrM fungi, a *T. calospora* and a *S. vermifera* isolate, has revealed an expanded family of Carbohydrate Active Enzymes (CAZymes) mainly involved in cellulose degradation (Kohler *et al*., 2015). This rich repertoire of plant cell wall degrading enzymes would suggest that rhizoctonia-like OrM fungi can spend part of their life cycle growing saprotrophically on plant litter in soil. However, their actual saprotrophic capabilities in nature are unclear and a limited saprotrophic potential is suggested, at least for some OrM fungi, by the results of metabarcoding investigations on the occurrence of rhizoctonia-like OrM fungi in the soil surrounding the roots of their orchid host. In particular, OTUs identified in the mycorrhizal roots of terrestrial orchids in Italy (Voyron *et al*., 2017), in Australia (Egidi *et al*., 2017), in the United States (Kaur *et al*., 2019) and in Denmark (Hartvig *et al*., 2024) could not be detected in the soil outside the orchid rhizosphere. As for *S. vermifera*, an experiment set up to evaluate fungal transmission to plants that can harbor this fungus as beneficial root endophyte (Ray *et al*., 2018) showed that transmission failed when the roots of different individuals were kept separate, thus indicating that migration of this fungus in root-free bulk soil is unlikely. Thus, the poor success of some OrM fungi in the “rhizoctonia” species complex as soil colonizers, as well as their potential behavior as root endophytes, raise intriguing questions on the actual niche of these fungi. Moreover, a recent study aimed at investigating the potential environmental reservoir of OrM fungi for *S. spiralis* (Calevo *et al*., 2021) showed that, despite the removal of environmental compartments (neighbouring plants and soil), a core of OrM fungi was able to colonize newly emergent roots, suggesting an internal transmission starting from either older roots or other orchid tissues. This finding further supports the hypothesis that some OrM fungi could be strictly associated to orchid tissues, spending most of their life cycle inside the plant and therefore qualifying as “ecologically obligate” orchid symbionts (Calevo *et al*., 2021).

Mycorrhizal fungi are reported to be strictly associated with the roots of land plants (Genre *et al*., 2020). However, given the peculiarities of OrM fungi in the “rhizoctonia” species complex, we have tested the hypothesis that at least some of these OrM fungi may use the whole plant as an ecological niche by colonizing not only the roots, but also other plant organs. Interestingly, rhizoctonia-like fungi were isolated from both roots and leaves of orchid species in the genus *Lepanthes* (Bayman *et al*., 1997) and in *Acampe praemorsa*, *Cymbidium aloifolium* and *Vanda testacea* (Behera *et al*., 2013). However, the phylogenetic position of these fungal isolates was not assessed, and it is therefore unclear whether roots and leaves hosted the same fungi. Here, we have used a metabarcoding approach to detect the occurrence of rhizoctonia-like taxa related to known OrM fungi in the mycobiota amplified from vegetative organs (roots, stems and leaves) of three terrestrial orchid species: *Spiranthes spiralis*, *Serapias vomeracea* and *Neottia ovata*. In one of the three orchid species, *S. spiralis*, we also investigated the seed mycobiota to test the possibility of a vertical transmission of OrM fungi.

## Material and Methods

### Orchid species

The plant species investigated in this work were *Spiranthes spiralis*, *Serapias vomeracea* and *Neottia ovata* (Fig. 1).

**Figure 1.**
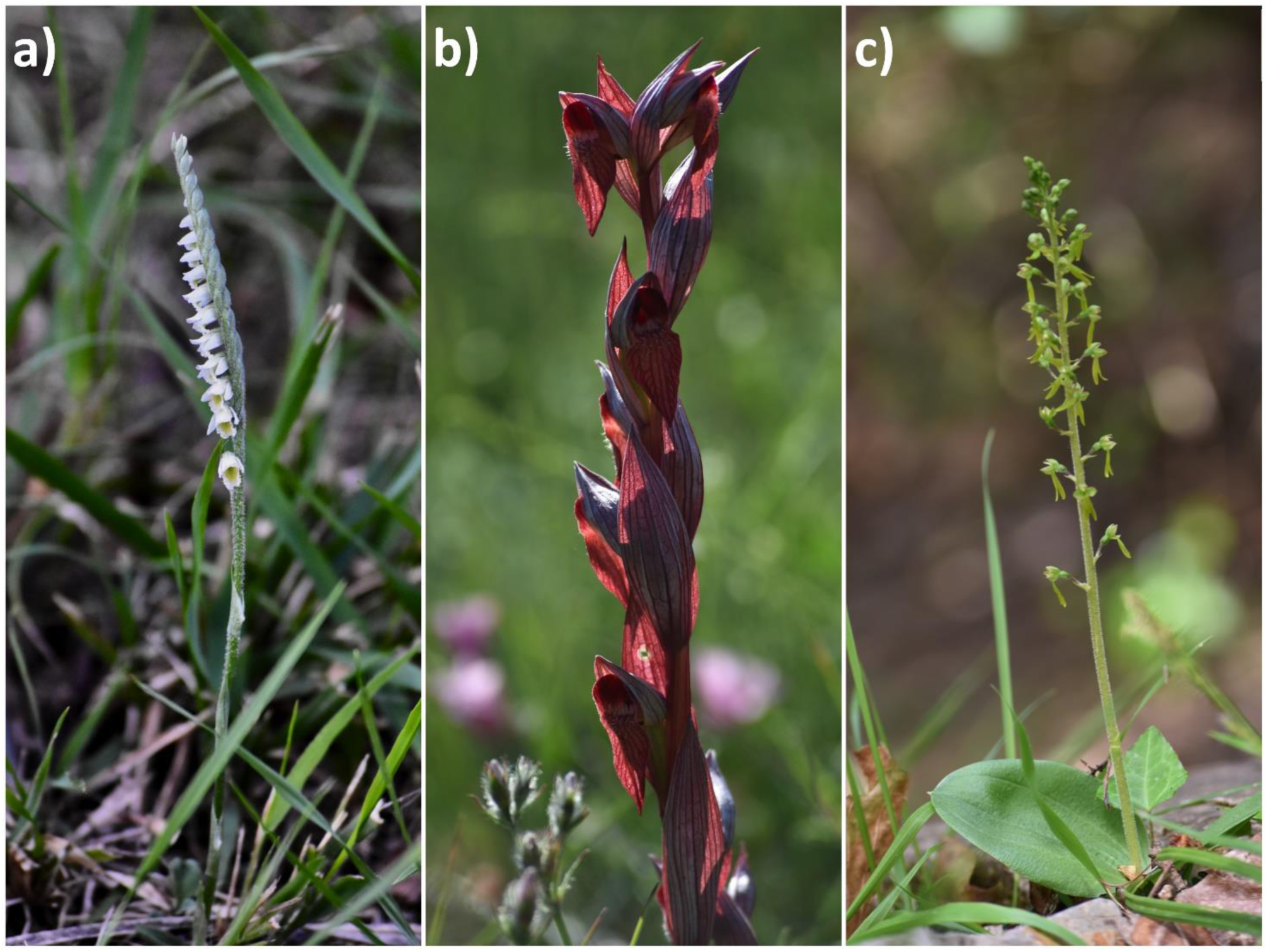
Floral morphology of the three target orchid species: (A) *Spiranthes spiralis*, (B) *Serapias vomeracea*, and (C) *Neottia ovata*. Each panel illustrates the distinct floral characteristics unique to each species, highlighting their morphological diversity.

*Spiranthes spiralis* (Subfamily Orchidoideae, Tribe Cranichideae) is a herbaceous orchid that flowers in late summer-autumn with a particular spiral-shaped inflorescence. It is widely distributed in Southern Europe and in the Mediterranean region, where it grows in pine, oak, chestnut, hornbeam and birch forests, dry meadows as well as in flat grasslands and semi-rocky areas. The preferred substrate is both calcareous and siliceous, with neutral pH. New leaves, formed at the same time or after the flower stem, stand together in a rosette beside the stem; the rhizome is periodically generated every year with new roots and stems (Arditti, 2002).

*Serapias vomeracea* (Subfamily Orchidoideae, Tribe Orchideae) is a bulbous herbaceous plant, with two underground globose rhizotubers and erect stems of purplish-vinous color, varying in height from 20 to 60 cm. The inflorescence, loose and elongated, is composed of a few spaced flowers. It can be found in sunny and wet meadows, on the edges of paths, in bushy environments from the plain up to 1200 m of altitude. It is the most widespread species in the genus *Serapias* and is distributed in most Europe (Arditti, 2002).

*Neottia ovata* (Subfamily Epidendroideae) is a perennial rhizomatous orchid regularly found in a wide range of habitats including woods, shrubs, hedges, calcareous pastures, dunes and marshes and, to a lesser extent, meadows. It grows on acid and calcareous substrates in damp and cold woods, mainly conifers, on sphagnum moss and carpets, often together with blueberry (*Vaccinium myrtillus*) from 900 to 2100 m a.s.l. The lower capsules in the inflorescence can mature and disperse the seeds even before the flowers placed higher in the inflorescence are pollinated. It is common throughout Europe (Arditti, 2002).

### Collection and Sterilization of Plant Samples

Various parts of the three orchid species were collected in the fall of 2016 and spring of 2017 from different locations within the provinces of Imperia (*Serapias vomeracea*), Savona (*Spiranthes spiralis*), and Genova (*Neottia ovata*) in Liguria, Italy. These parts included roots, stems, leaves, and seed pods. *Spiranthes spiralis* was collected twice: samples *S. spiralis*_1 were collected in 2016 (Calevo *et al*., 2021) and *S. spiralis*_2 were collected in 2017 to include capsules and seeds. Once collected, plant samples were immediately kept on ice within sterile containers and transported to the lab for storage at 4°C prior to examination. Sterilization of orchid tissues was performed as described in Alibrandi *et al*. (2021). Briefly, the above-ground parts of the plants underwent a sequential sterilization process involving a one-minute soak in 70% ethanol, a two-minute treatment with 2.5% sodium hypochlorite, and another one-minute immersion in 70% ethanol. This was followed by five washes in sterile water. The roots were first cleaned with sterile water, subjected to sonication, and then sterilized using 95% ethanol for 20 seconds and 5% sodium hypochlorite for three minutes, with seven subsequent rinses in sterile water. To verify the effectiveness of the sterilization, the final rinse water and imprints from the sterilized plant surfaces were cultured on various agar media(Luria Bertani - LB, King’s B) and monitored for microbial growth over a period of 4-7 days at 28°C.

### DNA Isolation, amplification and sequencing

Total DNA was isolated from approximately 100 mg of tissue, using the DNeasy Plant Mini Kit from QIAGEN, following the guidelines provided. DNA integrity and concentration were determined using a spectrophotometer (ND-1000 Spectrophotometer Nano DropH; Thermo Scientific, Wilmington, Germany). The ITS2 region of the nuclear ribosomal DNA was amplified from the extracted DNA through a semi-nested PCR method. Initially, the full ITS region (ITS1-5.8S-ITS2) was amplified using primer pairs specifically designed for orchid mycorrhizal fungi (Taylor & McCormick, 2008), ITS1-OFa, ITS1-OFb and ITS4-OF primers, and primers ITS1 and ITS4tul (Tul). A nested PCR was then conducted to amplify the ITS2 region using tagged primers ITS3mod and ITS4 (White *et al*., 1990). All amplifications were performed in three replicates. The PCR reactions were carried out with mixture and thermal cycling conditions as described in Calevo *et al*. (2020). The resulting PCR products were verified on a 1% agarose gel, replicates pooled together and purified with a Wizard SV Gel and PCR Clean-Up System (Promega), following the manufacturer’s instructions. Quantification was performed with QUBIT 2.0 (Thermo Fisher Scientific, Waltham, MA, USA) before preparing sequencing libraries, which were then sequenced using Illumina MiSeq technology (2 x 250 bp) by IGA Technology Services Srl (Udine, Italy).

### Sequence analyses

The initial step in analyzing the sequencing data involved merging the paired-end reads from each sample collection using PEAR v.0.9.2 (Zhang *et al*., 2014), setting a quality score cutoff at 28 for trimming and establishing a minimum read length of 200 base pairs post-trimming. The merged reads were then processed with the Quantitative Insights into Microbial Ecology (QIIME) v.1.8 software suite (Caporaso *et al*., 2010), adhering to specific criteria for sequence length, quality score, and primer mismatch tolerance as described in Voyron *et al*. (2017). Chimeric sequences were identified and excluded using an abundance-based approach with USEARCH61 (Edgar, 2010) within the QIIME framework. Clustering into operational taxonomic units (OTUs) was performed using a reference-guided method with a 98% similarity threshold, retaining only clusters with a minimum of 10 sequences. The UNITE database served as the reference for OTU identification and taxonomic classification (Abarenkov *et al*., 2010; Koljalg *et al*., 2013; http://unite.ut.ee, last accessed 25 May, 2019), employing the BLAST algorithm (Altschul *et al*., 1990) with a set e-value threshold of 1e-5. The most prevalent sequences within each OTU (the OTU representative sequences) of putative OrM fungi and fungi occurring in ≥80% plants in at least one orchid species were submitted to GenBank and recorded under the following string of accession numbers: PQ644909 - PQ645014.

We carried out the maximum likelihood (ML) analyses with representative sequences from ceratobasidioid, tulasnelloid and sebacinoid s.l. OTUs. Sequences that were the closest matches from BLAST searches, as well as sequences from a diverse range of terrestrial orchids, including the species under study, from various global locations and habitats, and sequences from other plants and fungal specimens were included in the analyses. The sequences were aligned using MAFFT (Katoh & Toh, 2010) with default settings, and the alignments were manually edited in MEGA v.7.0.26 (Kumar *et al*., 2016). ML estimations were conducted using RAxML v.8 (Stamatakis, 2014) over 1000 bootstrap replicates (Felsenstein, 1985), utilizing the GTR + GAMMA algorithm. The best tree topology was determined on the CIPRES Science Gateway (Miller *et al*., 2011); support values from bootstrapping runs were mapped on the globally best tree using the –f option of RAxML and -x 12345 as a random seed. Nodes with bootstrap support below 70% were not considered robust.

### Statistical analyses

#### Comparative Analysis of Root Sample Datasets

To facilitate comparative analysis across datasets derived from various tissue samples, including the newly collected *Spiranthes spiralis* and previously gathered tissues at the initial time point, a standardization process was implemented. This involved normalizing the sequencing depth to a consistent number of sequences (1000) for each sample, utilizing the *rarefy_even_depth* function within the ‘phyloseq’ package in R (R Core Team, 2021).

#### Analysis of Fungal Community Composition

The distribution of OTUs across different tissue categories was statistically compared using chi-squared tests to determine if there were significant differences in the proportions of OTUs recovered. The impact of orchid tissue on the composition of OrM fungal communities was investigated through permutational multivariate analysis of variance (PERMANOVA), employing 999 permutations within the adonis function of the ‘vegan’ package in R (Oksanen *et al*., 2013; R Core Team, 2021). Prior to this, the homogeneity of multivariate dispersions among groups was evaluated using the betadisper and permutest functions, also within the ‘vegan’ package (Oksanen *et al*., 2013). Constrained Analysis of Principal Coordinates (CAP) was also performed to evaluate the effects of species identity, organ type, and their interaction on OrM fungal community composition. CAP was chosen due to its robustness to heterogeneity in multivariate dispersions, as indicated by beta dispersion tests, which revealed significant differences in dispersion for organ type (P = 0.0382) but not for species (P = 0.1503).

Additionally, to visualize the distribution and relationships of OrM fungi, we employed R (R Core Team, 2021) packages such as ‘ggplot2’ (Wickham, 2016), ‘ggdendro’ (de Vries & Ripley, 2023), and ‘reshape2’ (Wickham, 2020) to create a heatmap of OrM fungi frequency in orchid tissues. This heatmap displayed hierarchical clustering of orchid samples based on the frequency patterns of OrM fungi association. To visualize OrM OTUs shared among different orchid tissues we plotted OTUs presence and frequency in chord diagrams generated using the ‘circlize’ package (Gu *et al*., 2014) for visualizing relationships, with layout adjustments made using the ‘gridExtra’ package (Auguie, 2017), and Venn diagrams.

## Results

After filtering and cleaning the Illumina MiSeq reads, we obtained 8,178,658 high-quality sequences that were clustered in 1353 OTUs (98% sequence identity). Only OTUs identified in the kingdom Fungi when blasted on the UNITE database and represented by at least 10 reads were considered for further analyses, for a total of 2,379,265 reads in 761 OTUs.

### The mycobiota in the vegetative organs of the three Mediterranean orchid species

The fungal OTUs retrieved from the vegetative organs of the three orchid species were mostly assigned to Ascomycota (471 OTUs, 51.4% of total OTUs), followed by Basidiomycota (297 OTUs, 32.4% of total OTUs) and unidentified OTUs (116 OTUs, 12.7% of total OTUs). Only very few OTUs were assigned to Mortierellomycota, Glomeromycota, Mucoromycota, Chytridiomycota, Kickxellomycota, Olpidiomycota and Rozellomycota, according to NCBI and UNITE taxonomic classification (each of the latter phyla accounting for <1.0% of total OTUs).

The mycobiota differed both among plant parts and among orchid species. In *S. spiralis*_1, *S. vomeracea* and *N. ovata*, the number of OTUs shared by at least two vegetative organs was significantly lower than the number of OTUs unique of a single organ (P < 0.00001, chi-squared tests). In all orchid species, in particular, roots and leaves in each plant shared a greater proportion of taxa with stems than with each other (Fig. S1). Generally, higher OTU richness was found in leaves than in roots (except in *S. spiralis*_2, in which OTU numbers did not differ significantly between leaves and roots; P < 0.00001, chi-squared tests). Despite the differences in OTU profiles between roots and leaves, most OTUs (394, 51.2% of total OTUs) were retrieved in both organs of the same individual orchid plant (70.1% of them being found in all vegetative organs of the same plant). Other OTUs (124 OTUs, 16.1%) were retrieved in either roots or leaves of different individuals of the same orchid species.

The different orchid species featured a different mycobiota. In fact, irrespective of the vegetative organ, a higher number of OTUs was exclusively found in a single orchid species (65.8-71.4% of total OTUs) than in more than one orchid species (P < 0.00001, chi-squared tests). In the PCA space, OTU profiles segregated based on orchid species rather than organ type (Fig. S2), with a partial overlap between the OTU profiles of *S. vomeracea* and *N. ovata*. A more distinct profile in both samplings characterized *S. spiralis*, especially when presence/absence was considered (Jaccard index).

Seventy-one OTUs (49.3% Ascomycota, 47.9% Basidiomycota, and one Mucoromycota) occurred in ≥80% samples of at least one organ in at least one orchid species (Table S1). Basidiomycota OTUs mostly included Cantharellales (35.3% of Basidiomycota OTUs, comprising several “rhizoctonias”), followed by Polyporales (15.2%) such as *Bjerkandera adusta*, *Daedaleopsis confragosa*, *Lentinus tuber-regium*, *Trametes versicolor*.

Clustering heatmap based on the dominant non-rhizoctonia OTUs revealed variable occurrence patterns of these OTUs in the different orchids (Fig. S3). Samples clustered again according to orchid species, OTU profiles in roots and stems being closely related in all orchids. Some taxa were retrieved in all vegetative organs of all orchids and included Cladosporiaceae (Ascomycota, Dothideomycetes, Capnodiales; OTU 127 *Mycosphaerella tassiana*, OTU 1180 *Cladosporium arthropodii*) and other Dothideomycetes (OTU 1105 *Aureobasidium pullulans*, OTU 11092 *Didymella exigua*, OTU 11092 *Pyrenochaetopsis leptospora*), as well as *Bjerkandera adusta* (OTU 733; Basidiomycota, Agaricomycetes, Polyporales, Meruliaceae). Other OTUs occurred in all organs of at least one orchid, while lacking in other species (e.g. OTUs 460 *Pholiotina brunnea* and 290 *Sistotrema brinkmannii* in *S. spiralis*_1, OTU 209 *Bifiguratus adelaidae* in *S. vomeracea* and *N. ovata*) (Fig.S3, Table S1).

### Potential OrM fungal taxa in *S. spiralis, S. vomeracea* and *N. ovata*

Previous investigations on *S. spiralis* (Tondello *et al*., 2012; Duffy *et al*., 2019; Calevo *et al*., 2021), *S. vomeracea* (Girlanda *et al*., 2011) and *N. ovata* (Jacquemyn *et al*., 2015; Těšitelová *et al*., 2015; Wang *et al*., 2023) revealed that rhizoctonia-like fungi that include tulasnelloid, ceratobasidioid and sebacinoid *s.l.* (sebacinoid *s.s*. and serendipitoid) fungi are the dominant OrM symbionts in these three terrestrial orchid species. We searched therefore, in the general database (Table S1), OTUs identified by Blast searches in the Cantharellales and Sebacinales, and selected 46 OTUs of potential OrM fungi belonging to tulasnelloid (11 OTUs), ceratobasidioid (27 OTUs) and sebacinoid *s.l.* fungi (9 OTUs). Representative sequences of these different OTUs were blasted against the NCBI nucleotide database. The same sequences were also aligned with reference sequences to better define their phylogenetic affinities, and the corresponding Maximum likelihood trees are shown in Figs. S4, S5 and S6 for tulasnelloid, ceratobasidioid and sebacinoid *s.l*. fungal OTUs, respectively.

With a single exception (OTU 184), the tulasnelloid OTUs clustered in a single clade (supported by 99% bootstrap) along with sequences previously obtained from the roots of *S. spiralis* (Calevo *et al*., 2021) and from other orchid species (i.e. from *Orchis sspp*., Cachapa Bailarote *et al*., 2012; Pecoraro *et al*., 2018; Calevo *et al*., 2020). Sequences in this tulasnelloid cluster were included in ‘species hypotheses’ of the UNITE database (Kõljalg *et al*., 2013) (SH1144039.08FU and SH1507552.08FU -DOI: SH1507552.08FU, Kõljalg *et al*., 2021) and assigned to *Tulasnella helicospora*. OTU 184 clustered separately, together with other sequences from *S. spiralis* (Duffy *et al*., 2019; Calevo *et al*., 2021) or other orchids (i.e., Liebel *et al*., 2010; Girlanda *et al*., 2011; Waud *et al*., 2014; Jacquemyn *et al*., 2015) as well as sequences from the soil of one of the study areas (Voyron *et al*., 2017; Fig. S4. Similarly, ceratobasidioid and sebacinoid s.l. OTUs were phylogenetically closely related to fungi identified in either the same or different orchid species, in roots of non-orchid plants or in the soil of one of the study areas (Figs. S5 and S6). Based on the clustering patterns in these phylogenetic analyses, the original 11 tulasnelloid, 26 ceratobasidioid and 9 sebacinoid s.l. OTUs were reduced respectively to 2, 23 and 8 OTUs for subsequent statistical analyses (see below).

### Many putative OrM fungi colonize the three orchid species systemically

Each plant hosted 10.9 ± 4.4 putative OrM fungal OTUs (mean ± standard deviation; 5-19 depending on the orchid individual). In each orchid species, ≥80% of plants hosted several ceratobasidioid and/or tulasnelloid fungi (Fig. 2, Table S1). All OTUs occurred in orchid roots (77.4% of them being retrieved in ≥60% plants in at least one orchid species), with the exception of the sebacinoid *s.l*. OTUs 1014 and 1154, which were only retrieved in stems of *S. vomeracea* and *S. spiralis*_2, respectively.

**Figure 2.**
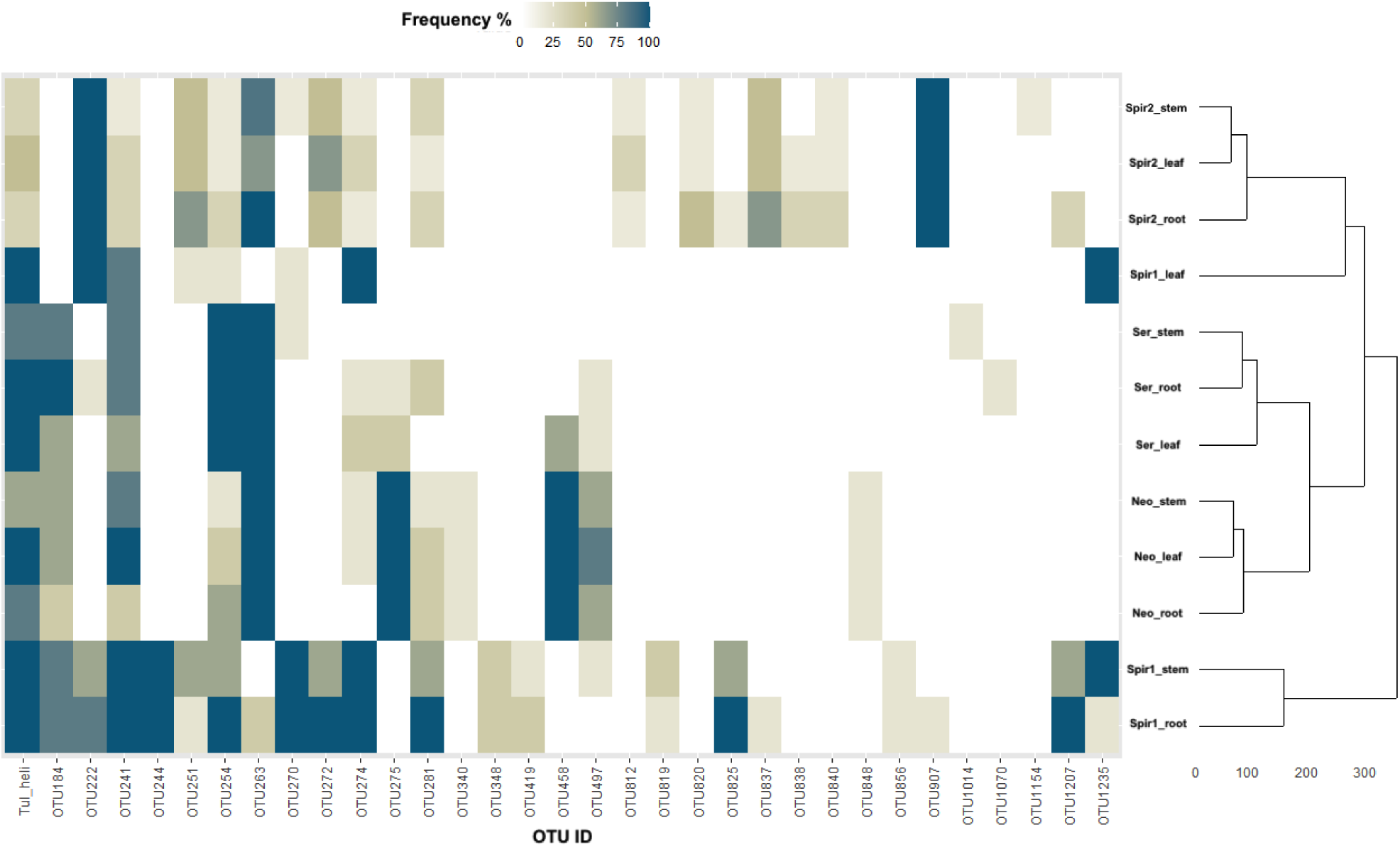
Heatmap and hierarchical clustering of OrM OTU frequency distribution across plant species and organs. The heatmap displays the frequency percentage (0-100%) of different OTUs in stem, leaf, and root samples from the target orchid species (*Spiranthes spiralis*_1*, Spiranthes spiralis*_2*, Serapias vomeracea, Neottia ovata*). The dendrogram on the right shows the hierarchical clustering of samples based on their OTU composition similarity. Color intensity from white to dark blue represents increasing OTU frequency.

To test our hypothesis that at least some OrM fungi may colonize their orchid host systemically, we traced the presence and distribution of the 33 OTUs assigned to sebacinoid *s.l*., tulasnelloid and ceratobasidioid fungi in the vegetative organs of the three orchid species. The results are summarized in the Venn diagrams in Fig. S7, that also reports the list of shared OTUs. A percentage of OTUs ranging from 16.7% in *S. vomeracea* to 50.0% in *N. ovata* was found in roots, leaves and stems, indicating that these fungi are able to colonize all the vegetative plant organs. In particular, a broad majority of the OrM fungi found in roots (84.8 %) was also retrieved from either stems or leaves in the same orchid plant, 61.5 % being found in all vegetative organs of the same plant. Although OrM fungi occurred more often in roots than in either stems or leaves (P = 0. 013131 and P < 0.00001, respectively), and more frequently in stems than in leaves (P = 0.004819, chi-squared tests; Fig. S8), they were detected more often in more than one organ than in a single one (P < 0.00001, chi-squared tests). Even more so, they were found more frequently in all three organs of the same plant than in any combination of two organs (P = 0.000027, P < 0.00001 and P < 0.00001 for the combinations roots&stems, roots&leaves and stems&leaves, respectively, chi-squared tests), or any single organ (P < 0.00001 for either roots, stems or leaves, chi-squared tests). Only the sebacinoid *s.l*. OTU 1070 was exclusively found in roots, but it was extremely sporadic, being retrieved from a single *S. vomeracea* plant.

The chord diagram in Fig. 3 shows, in the three orchid species, the distribution of tulasnelloid, ceratobasidioid and sebacinoid *s.l*. fungi in all organs of the same plant; for instance, the tulasnelloid OTU 162 assigned to *T. helicospora*, the ceratobasidioid OTUs 222, 254, 274_867, 263_319_1569, 275 and 907, and the sebacinoid *s.l*. OTU 458 were found in all organs of all individuals in at least one orchid species (Fig. 3 a-c). On the other hand, OTU distribution profiles could differ among organs (e.g. between leaves and either roots or stems in *S. spiralis*_2; P = 0. 00572 and P = 0. 0236, respectively, Kruskal-Wallis tests).

**Figure 3.**
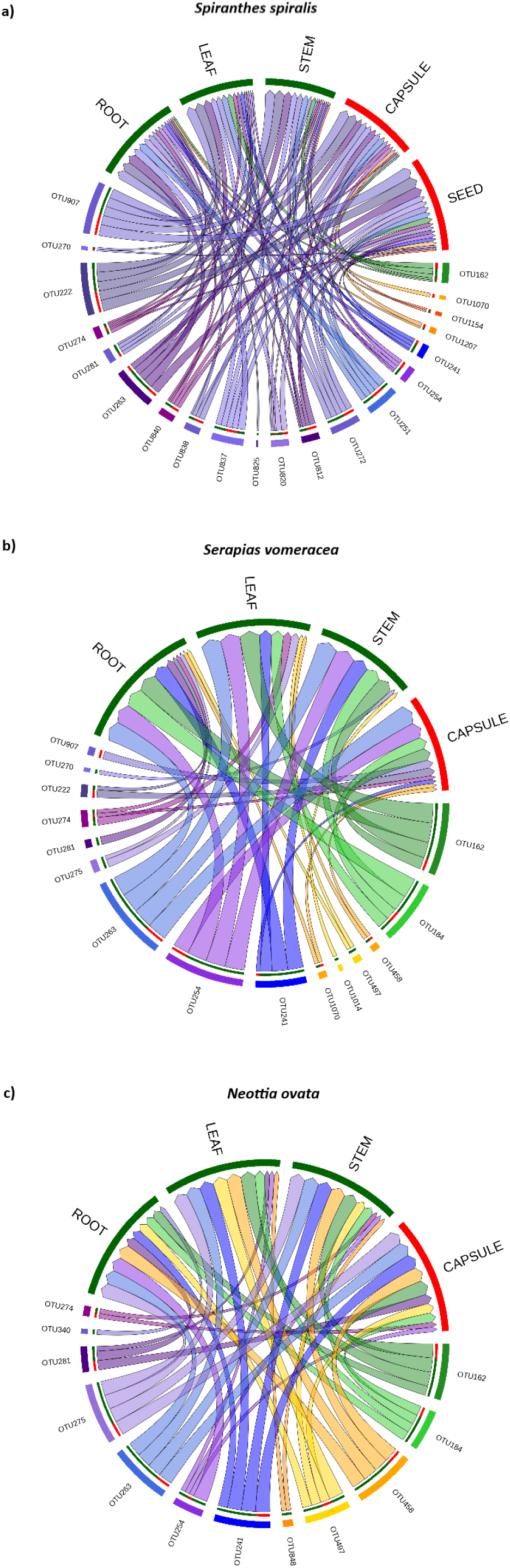
Circular diagrams illustrating the distribution of rhizoctonia-like OTUs (Operational Taxonomic Units) across different organs of the same plant for the three species: (A) *Spiranthes spiralis*_2, (B) *Serapias vomeracea*, (C) *Neottia ovata*. Each diagram shows connections between OTUs and plant organs (root, leaf, stem, capsule, seed), with color-coded segments representing different organs. The width of the connections indicates the abundance of OTUs in each organ, highlighting the diversity and distribution patterns within each species. Blue arches indicate ceratobasidioid OTUs, green arches tulasnelloid OTUs, while yellow arches indicate sebacinoid *s.l.* OTUs.

Like the general mycobiota, the OrM fungal assemblages in the different samples were significantly shaped by both the plant species and the plant organ (PERMANOVA, P < 0.001, Table S2). However, when subjected to NMDS analysis (3Dstress = 0.1298; linear fit, R2=0.81; Fig. 4), orchid samples mostly grouped according to orchid species rather than plant organ. In fact, different orchids featured different OrM fungi distribution profiles. For instance, *N. ovata* featured a lower proportion of occurrences only in roots, and a higher proportion of occurrences in leaves, than *S. spiralis*_1 (P = 0.048819 and P < 0.00001, respectively, chi-squared tests). *S. spiralis*_2 featured a lower number of OrM fungal OTUs per plant than *S. spiralis*_1 (P = 0.01762, Kruskall-Wallis test). OTUs which were exclusively found in a single orchid accounted for 41.9-58.1% of total OTUs in either roots, stems or leaves. The OrM fungal assemblage of *S. spiralis* in the first sampling (*S. spiralis*_1) appeared quite different from the profile of the second sampling (*S. spiralis*_2), mainly because of the higher frequency of tulasnelloid OTUs in the first sampling and the different distribution profiles of ceratobasidioid OTUs (see e.g. OTUs 241, 274 and 907, Fig. 2). Based on OTU frequencies, stem samples clustered with either root or leaf samples of the same orchid, while root and leaf samples never clustered directly together (Fig. 2).

**Figure 4.**
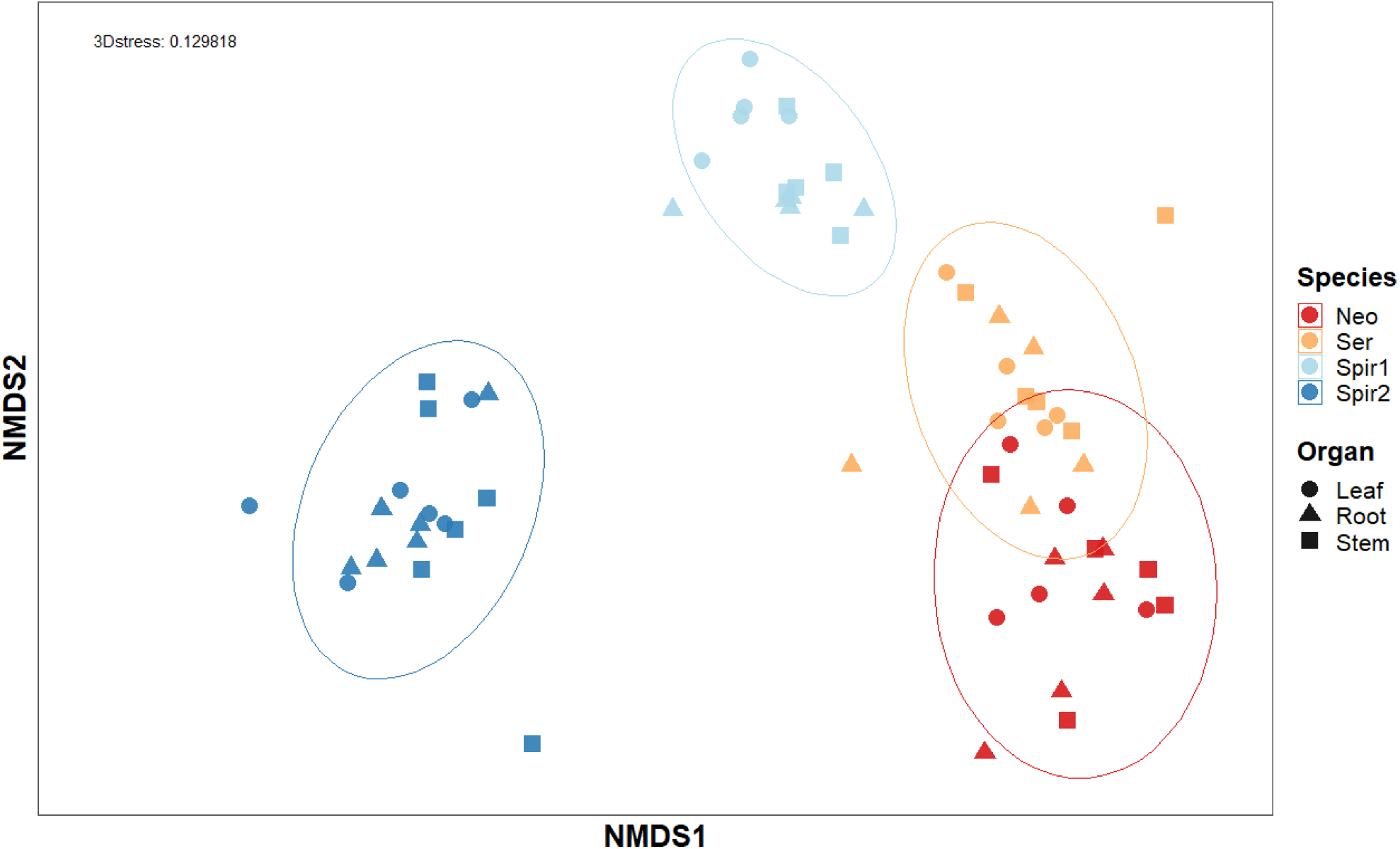
Non-metric multidimensional scaling (NMDS) plot of plant organ samples based on species. This NMDS plot visualizes the relationships among different plant organs (leaf, root, stem) based on OrM OTUs similarities across four species: *Neottia ovata, Serapias vomeracea, Spiranthes spiralis*_1, and *Spiranthes spiralis*_2. The axes represent NMDS1 and NMDS2 dimensions, with a stress value of 0.129818 indicating a good fit for the ordination. Data points are shaped according to organ type and colored by species.

### Colonization of reproductive organs by putative OrM fungi

The OrM fungal occurrence and distribution profiles were also assessed in the reproductive structures(capsules and/or seeds) of a subset of plants. Twenty-three tulasnelloid, ceratobasidioid and sebacinoid *s.l*. OTUs were retrieved from capsules/seeds of these plants, none of them being exclusively found in these reproductive structures (Fig. 3a-c). In fact, in each of the three orchid species, the OTUs retrieved from both vegetative organs and reproductive structures were found more often in more than one organ from the same plant than in capsules only (P = 0.001094, < 0.00001 and < 0.00001 for *S. vomeracea*, *N. ovata* or *S. spiralis*_2, respectively, chi-squared tests), or just in seeds in *S. spiralis*_2 (P < 0.00001, chi-squared tests). In the latter samples, OrM fungi occurred more frequently in both capsules and seeds than in either structure type alone (P < 0.00001 and 0.000639 for capsules and seeds, respectively, chi-squared tests). Only four OTUs (the two ceratobasidioid OTUs 340, Fig. 3b, and 825, Fig. 3c, and the sebacinoid *s.l*. OTUs 848, Fig.3b, and 1014, Fig.3a) were exclusively detected in the vegetative organs.

No significant difference was found in the distribution profiles of OrM fungal OTUs between vegetative and reproductive structures (CAP, P>0.05, Table S3), but a significant difference was found again between species, as well as for the interaction between species and organ type (CAP, P<0.001; Table S3), and samples grouped according to orchid species in the NMDS space (3Dstress = 0.1251; linear fit, R2=0.81; Fig. S9). For example, *T. helicospora* (OTU162), which occurred in most (≥60%) root, stem, leaf and capsule samples in either *S. vomeracea* or *N. ovata*, was sporadic in *S. spiralis*_2, while the sebacinoid *s.l*. OTUs 497, 516 and 458, which were not retrieved at all, or only rarely, in *S. spiralis*_2 and *S. vomeracea*, respectively, occurred in most *N. ovata* samples (Table S1).

## Discussion

The assemblages of fungi associated with internal orchid tissues have been investigated in several terrestrial and epiphytic species, where most studies focused on the root-associated mycobiota because of the major interest in OrM fungi (e.g. Bayman & Otero, 2006; Li *et al*., 2021; Ma *et al*., 2015; Rashmi *et al*., 2019; Sarsaiya *et al*., 2020; Bhatti & Thakur, 2022). Investigations on the endophytic mycobiota associated with other organs have instead been mostly restricted to economically important orchids, such as *Dendrobium* and *Vanilla* species (e.g. Yuan *et al*., 2009; Chen *et al.,* 2013, 2020; Parthibhan *et al*., 2017; Ma *et al*., 2022; Mosquera-Espinosa & Rodríguez-Mina, 2022; Parthibhan *et al*., 2017; Sarsaiya *et al*., 2020; Zhu *et al*., 2022).

### The mycobiota of vegetative plant organs include OrM fungi in all orchid species investigated

The primers used in this study retrieved large spectra of OTUs encompassing phylogenetically and ecologically diverse fungi, as found in other orchids (Li *et al*., 2021; Ma *et al*., 2015; Rashmi *et al*., 2019; Bhatti & Thakur, 2022; Cevallos *et al*., 2022). The dominant OTUs in the three orchid species investigated included fungi in the Ascomycota and Basidiomycota commonly reported in a wide range of hosts, such as species in *Alternaria*, *Aureobasidium*, *Cladosporium*, *Fusarium* (*Gibberella*), *Phoma sensu lato* (*Pyrenochaetopsis*, *Didymella*), *Malassezia* (Rashmi *et al*., 2019). Wood-decaying fungi (such as *Bjerkandera adusta* and other Polyporales, Table S1)found as endophytes of several tree and herbaceous species (Brum *et al*., 2012; Martin *et al*., 2015; Vaz *et al*., 2020; Faddetta *et al*., 2021; Parada *et al*., 2022), were also common.

As in other plants, the fungal assemblages were shaped by both the plant compartment and plant species (Harrison & Griffin, 2020; Trivedi *et al*., 2020). Indeed, each orchid species harbored a distinctive mycobiota and, within each orchid species, OTU profiles differed in distinct organs, suggesting they represent dissimilar microhabitats. Despite these differences, however, several members of the orchid mycobiota were shared by two or more organs, denoting either recruitment from a common environmental source, or the capacity for active fungal growth through different plant parts. In all orchid species, roots and leaves of the same plant exhibited greater overlap of taxa with stems than with each other, indicating the “intermediate” nature of stems as a fungal habitat along the vertical axis from roots to aboveground organs (Amend *et al*., 2019; Bahram *et al*., 2022). Profiles of the dominant OTUs were especially similar in roots and stems, which may be explained either by contamination of the lower parts of stems by rain splash from soil (Cooke & Rayner, 1984) or by upwards fungal growth from belowground organs.

Interestingly, 33 OTUs identified as putative OrM fungi in the “rhizoctonia” complex were found not only in roots but also in aerial organs of *S. spiralis*, *S. vomeracea* and *N. ovata*. Some of these OTUs (61.5%) were detected in roots, leaves and stems of the same plant, indicating systemic colonization of the internal orchid tissues. Like the general mycobiota, assemblages of these OrM fungi were shaped by both the orchid species host and by the plant organ.

### Role of rhizoctonia-like fungi in the orchid roots

As metabarcoding cannot test the actual mycorrhizal ability of the “rhizoctonia-like” fungi identified in the roots, such studies may potentially overestimate mycorrhizal fungal diversity (Albornoz *et al.,* 2021). However, the occurrence of hyphal pelotons, the typical OrM fungal structures, was microscopically verified in the roots of the three orchid species, and we assume that the peloton-forming fungi correspond to, or are included in, the spectrum of the tulasnelloid, ceratobasidioid and sebacinoid *s.l*. OTUs identified in the roots. The same OTUs clustered closely with sequences already reported as OrM in the same and in different orchid species, and a *Tulasnella helicospora* strain was recently demonstrated to induce symbiotic seed germination and protocorm colonization *in vitro* (Calevo *et al*., 2020). However, the possibility that some of the rhizoctonia-like fungi identified in our study do not form hyphal pelotons in the orchid roots cannot be ruled out entirely and is discussed later.

The role of OrM fungi in the uptake and transfer of nutrients to adult orchids has been demonstrated *in vitro* (Cameron *et al*., 2006, 2007, 2008; Read *et al*., 2024) and in nature (Gomes *et al*., 2023; Zahn *et al*., 2023, 2024). Nutrient transfer to the host plant is thought to occur across the plant-fungus interface that surrounds the intracellular hyphal pelotons formed in OrM protocorms and roots (Perotto & Balestrini, 2024). Several questions remain open on the nature of nutrients transferred to the orchid host by rhizoctonia-like OrM fungi (Fochi *et al*., 2017; De Rose *et al*., 2023), but the role of these fungi in the supply of nutrients to the orchid host and the site of nutrient transfer in mycorrhizal roots (i.e., the hyphal pelotons) have never been questioned.

### Potential role of rhizoctonia-like fungi as endophytes in orchid roots and aerial parts

Although the nutritional benefits for the host plant are generally ascribed to mycorrhizal fungi (Selosse *et al*., 2022), there is growing evidence that specialized structures such as those formed in mycorrhizal associations are not essential for improved nutrient uptake and plant growth promotion, as some root-associated non-mycorrhizal fungal endophytes can aid nutrient absorption and nutrient allocation to different plant parts through a range of mechanisms (Poveda *et al*., 2021; Sarkar *et al*., 2021; Verma *et al*., 2021; Watts *et al*., 2023). Some fungal endophytes can in fact increase the elemental composition (especially nitrogen and phosphorous) in roots and shoots of inoculated plants, *S. vermifera* being the best studied example in non-orchid hosts (Verma *et al*., 2021). Therefore, we cannot exclude an additional role in plant nutrition by other fungi among those identified in the orchid root mycobiota, or by rhizoctonia-like fungi that may possibly colonize the orchid root without forming pelotons.

The type of interaction with the host plant and the role of these rhizoctonia-like OrM fungi in the vegetative aerial plant parts (stems and leaves) is currently unknown, and we can only make speculations on their functional role based on research on other fungal endophytes. The contribution of some fungal endophytes which are not (or not exclusively) root endophytes to plant nutrient acquisition has been reported (Sarkar *et al*., 2021). For instance, *Epichloë* species (Clavicipitaceae, Ascomycota) are endophytic symbionts of cool-season grasses (White, 1987). Whereas the *Epichloë* mycelium is found throughout the above-ground portions of the grass plant, roots are typically not colonized (Rodriguez *et al*., 2009; Mosaddeghi *et al*., 2021). These endophytes increase plant nutrient uptake by multiple mechanisms, including enhanced root proliferation by indole-3-acetic acid production (Mosaddeghi *et al*., 2021) and release of root exudates that facilitate solubilization and absorption of some nutrients, such as P, K and Fe (Malinowski & Belesky, 2000; Soto-Barajas *et al*., 2016).

These findings indicate that connection with the soil environment is not an essential requirement for aid in plant nutrient uptake by fungi associated with aerial plant parts (Kariman *et al*., 2018). For fungi establishing systemic infection of orchids, like the rhizoctonia-like fungi identified in this study, the hyphal connection between roots and aerial plant parts may represent an additional route for nutrient allocation in the host plant. Moreover, non-nutritional benefits also accrue to endophyte-infected plant hosts. Colonization by *Epichloë* spp. also confers plant tolerance to individual or combined abiotic (i.e., drought, salinity, mineral and temperature extremes), as well as biotic stresses, including local and systemic resistance to pathogens (Mosaddeghi *et al*., 2021). Intriguingly, some *Ceratobasidium* isolates from OrM orchid roots were effective as biocontrol agents of *Rhizoctonia solani* (= *Thanatephorus cucumeris*), a pathogen that causes sheath blight in rice (Mosquera-Espinosa *et al*., 2013). Other OrM isolates (one *Ceratobasidium* sp. and two *Tulasnella* spp. isolates) showed some antagonistic *in vitro* abilities against *Fusarium oxysporum* f.sp. *vanillae*, the causative agent of *Fusarium* wilt in the orchid *Vanilla planifolia*, but no biocontrol activity when inoculated *in vivo* (Manrique-Barros *et al*., 2023).

Thus, although their role requires further investigations, OrM fungi have multiple opportunities, by virtue of their systemic colonization of orchid tissues, to benefit their plant hosts at the adult stage.

### Potential vertical transmission of OrM rhizoctonia-like fungi and benefits for the plant

Whatever their structural/functional relationship with the orchid tissues, the capacity of OrM fungi for systemic colonization of the orchid organs has implications for the persistence and spread of these fungi in the environment. The identification, in the capsules of the three orchid species, of a core mycobiota including rhizoctonia-like OrM fungi also found in vegetative organs, indicates that these fungi could also colonize the flower and the reproductive organs. For *S. spiralis*, it was possible to track putative OrM fungal OTUs also in the seeds released from mature capsules. Similarly, a metabarcoding study by Wang *et al*. (2022) retrieved the same tulasnelloid OTU in mature mycorrhizal roots and in capsules of wild *Cymbidium goeringii* plants in China. Although more samples and species need to be investigated, these findings open the intriguing possibility that some OrM fungi could be vertically transmitted.

Vertical transmission of systemic fungal endophytes from a mother plant to the new seedlings *via* the seeds has been originally reported for *Epichloë* species in grasses (Liu *et al*., 2017), but it is likely more widespread and includes many more species than previously thought (Hodgson *et al*., 2014; Shahzad *et al*., 2018; Abdelfattah *et al*., 2021; Özkurt *et al*., 2020; Fort *et al*., 2021). Given the importance of OrM fungi in orchid seed germination and recruitment, a strategy that allows the minute orchid seeds to travel with their own symbiotic fungi would likely be evolutionarily successful, provided that the same fungi colonizing adult plants can also promote seed germination and protocorm development. The spectrum of OrM fungi may in fact change during protocorm development into adult plants in some orchid species (Rasmussen *et al*., 2015; Ventre Lespiaucq *et al*., 2021), but most OrM fungi able to promote symbiotic seed germination were originally isolated from adult plants (e,g, Fuji *et al*., 2020; McCormick *et al*., 2021; Těšitelová *et al*., 2022; Calevo & Duffy, 2023; Fernández *et al*., 2023; Chamara *et al*., 2024; Freestone *et al*., 2024). Moreover, most results on the local restriction of orchid distribution by OrM fungal abundance demonstrate that seed germination is higher near adult orchids (see in McCormick *et al*., 2018), suggesting at least partially shared OrM fungi.

### Possible ecological benefits of systemic endophytism for OrM fungi

For plants such as orchids, that rely heavily on symbiotic fungi throughout their life history stages, the possibility to host important fungal taxa as systemic colonizers and to transmit them vertically to the offspring *via* the seeds could be a successful strategy for survival and dispersal (Khan *et al*., 2015; Shahzad *et al*., 2018). In return, OrM fungi could use their host plants for survival and persistence in the environment (Selosse, 2014; Selosse & Martos, 2014; Oja *et al*., 2015; McCormick *et al*., 2016; Voyron *et al*., 2017). Studies by McCormick *et al*. (2018) and Waud *et al*. (2016) showed that OrM fungi were abundant in small soil patches closely tied to the locations of adult orchids, thus suggesting that adult orchid plants provide their OrM fungal symbionts with a ‘refuge’. Under natural conditions, the prevalence of some rhizoctonia-like OrM fungi inside the host plant and their undetectability in soils (Voyron *et al*., 2017; Egidi *et al*., 2017; Kaur *et al*., 2019; Hartvig *et al*., 2024) further suggest that some symbiotic fungi may even be unable to grow outside the host plant. For such fungi, specialization on systemic biotrophism may lead to an ecological advantage, as plants may represent a refugium from the stress and competition conditions associated with saprotrophic growth in soil (Cooke & Rayner, 1984).

Endophytism appears to be often associated with environmental abiotic stress, like heat stress (Matheny *et al*., 2018; Raudabaugh *et al*., 2020; Fox *et al*., 2022). In Mediterranean environments, like those where our orchid species grow, foliar endophytism may be a possible adaptation to cope not only with heath, but also with moisture limitation (Vaz *et al*., 2020). Thus, although questions remain open on the actual capabilities of OrM fungi to grow in the soil as saprotrophs, endophytism may have been an alternative for these fungi, especially in stressful environments such as the Mediterranean region.

## Conclusions

Our results show that rhizoctonia-like fungi in the tulasnelloid, ceratobasidioid and sebacinoid *s.l*. clades not only associate with orchid roots, where they likely form OrM, but they are also capable of endophytic growth in the aerial tissues of their orchid hosts, including reproductive organs. Further investigations are required to assess the colonization pattern, the viability and the functional role of rhizoctonia-like OrM fungi in the aerial plant organs and in seeds, but these first results may suggest a specific strategy for survival and dispersal that would benefit both the orchid host and the associated fungal partners.

An intriguing question is the evolution of the interactions between rhizoctonia-like OrM fungi and their orchid host. As many rhizoctonia-like OrM fungi are also endophytes in non-orchid plants, it has been hypothesized that extant OrM fungi were recruited among root endophytic fungi that colonized orchid ancestors (the ‘waiting room’ hypothesis, Selosse *et al*., 2022). Our findings that OrM fungi can colonize at the same time both roots and aerial plant organs of their extant orchid hosts make the scenario far more complex and expand the life history traits and potential ecological roles of OrM fungi.

## Acknowledgments

J.C. was funded by a European Commission Marie Skłodowska-Curie Global Fellowship (grant agreement No. 101031324 “FORECAST – Quantifying the impact of climate change on orchid mycorrhizal symbiosis in Mediterranean biodiversity hotspots”). P.A. was supported by a PhD fellowship by MUR. Research was supported by National funds PRIN 2017TP3SJL “Mediterranean orchids as a model system to quantify intrinsic and extrinsic factors influencing distribution and conservation-oriented restoration”.

## Competing interests

Authors declare no competing interests

## Author contributions

All authors contributed to the study conception and design. Material preparation and data collection were performed by J.C. and P.A. Data analysis was performed by all authors. The first draft of the manuscript was written by S.P., J.C. and M.G., and all authors commented on previous versions of the manuscript, read and approved the final manuscript.

## Data availability

We confirm that, should the manuscript be accepted, the data supporting the results will be archived in Zenodo under the DOI: 10.5281/zenodo.14223971. The ITS representative sequences of fungi from Supplementary Table S1 were submitted to GenBank and recorded under the following string of accession numbers: PQ644909 - PQ645014.

## Supplementary material

**Table S1**. Taxonomic affiliation (according to NCBI) of the OTUs occurring in ≥80% plants in at least one orchid species and the 33 rhizoctonia-like OTUs. Occurrences in the orchid vegetative organs (percentage of plants hosting in each vegetative organ in each orchid species) are also reported. Spir1= *Spiranthes spiralis*_1; Spir2= *S. spiralis*_2; Ser= *Serapias vomeacea*; Neo= *Neottia ovata*.

**Table S2**. PERMANOVA results assessing the effects of species, organ, and their interaction on the Bray - Curtis and Jaccard dissimilarity matrixes of the OrM fungi dataset. The analysis was conducted using 9999 permutations to determine the significance of each term, revealing that species, organ, and their interaction significantly influence the community composition, as indicated by the F-values and R² values.

**Table S3**. Summary of Constrained Analysis of Principal Coordinates (CAP) results comparing the effects of species identity, organ type (vegetative or reproductive), and their interaction on OrM fungal community composition. The table shows the F-statistic, R² (proportion of variance explained), p-values, and significance levels for each factor and their interaction. Analyses were performed using both Bray-Curtis and Jaccard distance metrics. Significance levels are indicated as: *** (p < 0.001), ** (p < 0.01), * (p < 0.05), ns (not significant). The results show that species identity and the interaction between species and organ type have a highly significant effect on OrM fungal community composition (p < 0.001) with both distance metrics.

**Figure S1**. Venn diagrams showing the number of shared and organ-specific fungal OTUs in the roots, stems and leaves of the same plant for each of the three orchid species: *Spiranthes spiralis*, *Serapias vomeracea* and *Neottia ovata*. For *S. spiralis*, plants were sampled in two consecutive years.

**Figure S2**. Principal Component Analysis (PCA) of fungal communities associated with different orchid organs based on Jaccard (left) and Bray-Curtis (right) dissimilarity matrices. The analysis includes fungal OTUs from three orchid taxa (*Neottia*, *Serapias*, and two *Spiranthes* group samples) and three plant organs (leaf, root, and stem). Points are shaped according to the organ type (circles = leaf, triangles = root, crosses = stem) and colored by orchid taxa. The percentage of variance explained by each principal component is shown on the axes. PCoA1 explains 15.33% and 18.25% of the total variance for Jaccard and Bray-Curtis analyses respectively.

**Figure S3**. Heatmap and hierarchical clustering of the dominant non-OrM fungal OTUs across plant species and organs. The heatmap displays OTU frequency percentages (0-100%) in different plant organs (stem, root, leaf) of the target orchids: *Spiranthes spiralis_1* (Spir1)*, Spiranthes spiralis_2* (Spir2)*, Neottia ovata* (Neo)*, Serapias vomeracea* (Ser). Hierarchical clustering (dendrogram) shows sample relationships. Color intensity from white to dark blue represents increasing OTU frequency.

**Figure S4**. Maximum likelihood tree obtained from the ITS2 sequence alignment of tulasnelloid fungi. *Multicalvula vernalis* was used as outgroup. Bootstrap support values above 70% (1000 maximum likelihood replicates) are reported. OTUs found in this study are indicated in bald, sequences retrieved in previous studies from the same orchid species are in red.

**Figure S5**. Maximum likelihood tree obtained from the ITS2 sequence alignment of ceratobasidioid fungi. *Tricholoma portentosum* and *Laccaria bicolor* were used as outgroup taxa. Bootstrap support values above 70% (1000 maximum likelihood replicates) are reported. OTUs found in this study are indicated in bald, sequences retrieved in previous studies from the same orchid species are in red.

**Figure S6**. Maximum likelihood tree obtained from the ITS2 sequence alignment of fungi assigned to Sebacinales. *Tremiscus helvelloides* was used as an outgroup taxon. Bootstrap support values above 70% (1000 maximum likelihood replicates) are reported. OTUs found in this study are indicated in bald, sequences retrieved in previous studies from the same orchid species are in red.

**Figure S7**. Venn diagrams showing the number of tulasnelloid, ceratobasidioid and sebacinoid s.l. OTUs in the roots, stems and leaves of the same plant for each of the three orchid species: *Spiranthes spiralis*, *Serapias vomeracea* and *Neottia ovata*. The list of OTUs shared between different organs is provided. For *S. spiralis*, plants were sampled in two consecutive years.

**Figure S8**. The larger pie charts depict the share of occurrences of the 33 rhizoctonia OTUs in either all (RSL), two (RS, roots&stem; RL, roots&leaves and LS, leaves&stem), or a single vegetative organ (R, roots; S, stem; L, leaves) in the same plant for each orchid species. The smaller charts summarize the share of total occurrences in roots, stems and leaves.

**Figure S9**. Non-metric multidimensional scaling (NMDS) plot illustrating the relationships among different plant organs and species based on their characteristics. The axes represent NMDS1 and NMDS2 dimensions. Data points are shaped according to organ type (capsule, leaf, root, seed, stem) and colored by species (*Neottia ovata*, *Serapias vomeracea*, *Spiranthes spiralis*_2). Ellipses indicate clustering patterns, with a stress value of 0.125118, suggesting a good fit for the ordination.

## Notes

### Competing Interest Statement

The authors have declared no competing interest.

