## Supplementary Information for "Breaking boundaries: fungi in the “rhizoctonia” species complex exhibit systemic colonization in three terrestrial orchid species"

**Table S1**. Taxonomic affiliation (according to NCBI) of the OTUs occurring in ≥80% plants in at least one orchid species and the 33 rhizoctonia-like OTUs. Occurrences in the orchid vegetative organs (percentage of plants hosting in each vegetative organ in each orchid species) are also reported. Spir1= *Spiranthes spiralis*_1; Spir2= *S. spiralis*_2; Ser= *Serapias vomeacea*; Neo= *Neottia ovata*.

| OTU no. | Taxonomic affiliation | | | | | Percentage of hosting plants | | | | | | | | | | | |
|---|---|---|---|---|---|---|---|---|---|---|---|---|---|---|---|---|---|
| | Species | Phylum | Class | Order | Family | Spir1 | Spir1 | Spir1 | Spir2 | Spir2 | Spir2 | Ser | Ser | Ser | Neo | Neo | Neo |
| | | | | | | Roots | Stem | Leaves | Roots | Stem | Leaves | Roots | Stem | Leaves | Roots | Stem | Leaves |
| **222** | **Ceratobasidioid sp.** | Basidiomycota | Agaricomycetes | Cantharellales | Ceratobasidiaceae | 13 | | 6 | 67 | 67 | 83 | 20 | | 13 | | | 13 |
| **241** | **Ceratobasidioid sp.** | Basidiomycota | Agaricomycetes | Cantharellales | Ceratobasidiaceae | 87 | 40 | 44 | | | | 30 | 71 | 50 | 33 | 67 | 75 |
| **244** | **Ceratobasidioid sp.** | Basidiomycota | Agaricomycetes | Cantharellales | Ceratobasidiaceae | 93 | 40 | | | | | | | | | | |
| **251** | **Ceratobasidioid sp.** | Basidiomycota | Agaricomycetes | Cantharellales | Ceratobasidiaceae | 7 | | | 33 | 50 | 50 | | | | | | 13 |
| **254** | **Ceratobasidioid sp.** | Basidiomycota | Agaricomycetes | Cantharellales | Ceratobasidiaceae | 11 | 40 | | 17 | | | 70 | 100 | 38 | 22 | | 13 |
| **263** | **Ceratobasidioid sp.** | Basidiomycota | Agaricomycetes | Cantharellales | Ceratobasidiaceae | | | | 17 | 17 | 17 | 100 | 100 | 100 | 89 | 100 | 75 |
| **270** | **Ceratobasidioid sp.** | Basidiomycota | Agaricomycetes | Cantharellales | Ceratobasidiaceae | 93 | 60 | | | | | | | | | | |
| **272** | **Ceratobasidioid sp.** | Basidiomycota | Agaricomycetes | Cantharellales | Ceratobasidiaceae | 100 | 40 | | 50 | 50 | 50 | | | | | | 13 |
| **274** | **Ceratobasidioid sp.** | Basidiomycota | Agaricomycetes | Cantharellales | Ceratobasidiaceae | 93 | 80 | 44 | | 17 | 17 | 10 | 14 | 25 | | 17 | 13 |
| **275** | **Ceratobasidioid sp.** | Basidiomycota | Agaricomycetes | Cantharellales | Ceratobasidiaceae | | | | | | | | | 25 | 89 | 100 | 100 |
| **281** | **Ceratobasidioid sp.** | Basidiomycota | Agaricomycetes | Cantharellales | Ceratobasidiaceae | 100 | 40 | | | 17 | | 30 | 14 | 13 | 11 | 33 | 25 |
| **340** | **Ceratobasidioid sp.** | Basidiomycota | Agaricomycetes | Cantharellales | Ceratobasidiaceae | | | | | | | | | | 11 | | 13 |
| **348** | **Ceratobasidioid sp.** | Basidiomycota | Agaricomycetes | Cantharellales | Ceratobasidiaceae | 7 | | | | | | | | | | | |
| **812** | **Ceratobasidioid sp.** | Basidiomycota | Agaricomycetes | Cantharellales | Ceratobasidiaceae | | | | 17 | | | | | | | | |
| **819** | **Ceratobasidioid sp.** | Basidiomycota | Agaricomycetes | Cantharellales | Ceratobasidiaceae | 13 | | | | | | | | | | | |
| **820** | **Ceratobasidioid sp.** | Basidiomycota | Agaricomycetes | Cantharellales | Ceratobasidiaceae | | | | 17 | | | | | | | | |
| **825** | **Ceratobasidioid sp.** | Basidiomycota | Agaricomycetes | Cantharellales | Ceratobasidiaceae | 53 | | | | | | | | | | | |
| **837** | **Ceratobasidioid sp.** | Basidiomycota | Agaricomycetes | Cantharellales | Ceratobasidiaceae | 7 | | | 33 | 17 | 33 | | | | | | 13 |
| **838** | **Ceratobasidioid sp.** | Basidiomycota | Agaricomycetes | Cantharellales | Ceratobasidiaceae | | | | 17 | | 17 | | | | | | |
| **840** | **Ceratobasidioid sp.** | Basidiomycota | Agaricomycetes | Cantharellales | Ceratobasidiaceae | | | | 17 | | | | | | | | |
| **856** | **Ceratobasidioid sp.** | Basidiomycota | Agaricomycetes | Cantharellales | Ceratobasidiaceae | | | | | | | | | | | | |
| **907** | **Ceratobasidioid sp.** | Basidiomycota | Agaricomycetes | Cantharellales | Ceratobasidiaceae | | | | 50 | 67 | 83 | | | | | | |
| **419** | ***Thanatephorus*** sp. | Basidiomycota | Agaricomycetes | Cantharellales | Ceratobasidiaceae | 7 | | | | | | | | | | | |

| | | | | | | | | | | | | | | | | |
|---|---|---|---|---|---|---|---|---|---|---|---|---|---|---|---|---|
| **162** | ***Tulasnella helicospora*** | Basidiomycota | Agaricomycetes | Cantharellales | Tulasnellaceae | 100 | 100 | 100 | | | | 70 | 71 | 75 | 67 | 50 | 100 |
| **184** | **Tulasnelloid** sp. | Basidiomycota | Agaricomycetes | Cantharellales | Tulasnellaceae | 7 | 20 | | | | | 30 | 43 | 50 | 33 | 67 | 50 |
| **458** | **Sebacinoid sp.** | Basidiomycota | Agaricomycetes | Sebacinales | | | | | | | | | | 13 | 100 | 100 | 88 |
| **497** | **Sebacinoid sp.** | Basidiomycota | Agaricomycetes | Sebacinales | | | | | | | | | | 13 | 56 | 33 | 63 |
| **848** | **Sebacinoid sp.** | Basidiomycota | Agaricomycetes | Sebacinales | | | | | | | | | | | | | 13 |
| **1014** | **Sebacinoid sp.** | Basidiomycota | Agaricomycetes | Sebacinales | | | | | | | | | 14 | | | | |
| **1070** | **Sebacinoid sp.** | Basidiomycota | Agaricomycetes | Sebacinales | | | | | | | | 10 | | | | | |
| **1154** | **Sebacinoid sp.** | Basidiomycota | Agaricomycetes | Sebacinales | | | | | | | | | | | | | |
| **1207** | **Sebacinoid sp.** | Basidiomycota | Agaricomycetes | Sebacinales | | 20 | 20 | | | | | | | | | | |
| **1235** | **Sebacinoid sp.** | Basidiomycota | Agaricomycetes | Sebacinales | | 7 | | 6 | | | | | | | | | |
| **1230** | ***Russulales*** sp. | Basidiomycota | Agaricomycetes | Russulales | | | 20 | 6 | 50 | 17 | 83 | 30 | 29 | 25 | 44 | 50 | 50 |
| **1105** | ***Aureobasidium pullulans*** | Ascomycota | Dothideomycetes | Dothideales | Dothioraceae | 47 | 100 | 69 | 67 | 67 | 83 | 20 | 57 | 63 | 44 | 67 | 88 |
| **1149** | ***Xylomyces*** sp. | Ascomycota | Dothideomycetes | Jahnulales | Aliquandostipitaceae | 100 | 100 | 94 | | | | | 29 | 13 | | 17 | 50 |
| **1318** | ***Mycosphaerellaceae* sp.** | Ascomycota | Dothideomycetes | Mycosphaerellales | Mycosphaerellaceae | 7 | | | 67 | 50 | 50 | 20 | 43 | 13 | 100 | 100 | 100 |
| **1450** | ***Lophium arboricola*** | Ascomycota | Dothideomycetes | Mytilinidiales | Mytilinidiaceae | 93 | 40 | 56 | | | 17 | | 29 | | | | |
| **722** | ***Pyrenochaetopsis leptospora*** | Ascomycota | Dothideomycetes | Pleosporales | Cucurbitariaceae | 7 | 100 | 94 | | 17 | 83 | 40 | 86 | 88 | 67 | 83 | 100 |
| **1092** | ***Didymella exigua*** | Ascomycota | Dothideomycetes | Pleosporales | Didymellaceae | 87 | 100 | 100 | 100 | 100 | 100 | 60 | 86 | 63 | 56 | 100 | 75 |
| **1544** | ***Paraphaeosphaeria parmeliae*** | Ascomycota | Dothideomycetes | Pleosporales | Didymosphaeriaceae | 13 | 20 | | | | | | | | | | |
| **1542** | ***Plenodomus biglobosus*** | Ascomycota | Dothideomycetes | Pleosporales | Leptosphaeriaceae | 100 | 100 | 100 | | | 17 | | 29 | 13 | | | |
| **1272** | ***Herpotrichia juniperi*** | Ascomycota | Dothideomycetes | Pleosporales | Melanommataceae | 100 | 80 | 94 | | | | | 29 | 13 | 22 | 17 | 13 |
| **1102** | ***Acrocalymma vagum*** | Ascomycota | Dothideomycetes | Pleosporales | Morosphaeriaceae | 20 | | | 83 | 100 | 100 | 10 | | | | | 13 |
| **1255** | ***Septoria oenanthicola*** | Ascomycota | Dothideomycetes | Pleosporales | Mycosphaerellaceae | 20 | 40 | 6 | 50 | 33 | 50 | 90 | 100 | 100 | 78 | 100 | 88 |
| **1238** | ***Sphaerulina pseudovirgaureae*** | Ascomycota | Dothideomycetes | Pleosporales | Mycosphaerellaceae | | 40 | 94 | | | | 60 | 100 | 100 | 56 | 83 | 88 |
| **1027** | ***Mycosphaerellaceae*** sp. | Ascomycota | Dothideomycetes | Pleosporales | Mycosphaerellaceae | 100 | 80 | 94 | | | | 30 | 29 | 25 | | 17 | 38 |
| **1459** | ***Periconia digitata*** | Ascomycota | Dothideomycetes | Pleosporales | Periconiaceae | 7 | 20 | | 67 | 83 | 83 | | | | | | 13 |
| **1121** | ***Alternaria alternata*** | Ascomycota | Dothideomycetes | Pleosporales | Pleosporaceae | 100 | 100 | 100 | 100 | 100 | 100 | 70 | 100 | 88 | 67 | 67 | 88 |
| **1458** | ***Alternaria metachromatica*** | Ascomycota | Dothideomycetes | Pleosporales | Pleosporaceae | 7 | 60 | 25 | 100 | 100 | 100 | 40 | 14 | 75 | | | 25 |
| **1049** | ***Stemphylium vesicarium*** | Ascomycota | Dothideomycetes | Pleosporales | Pleosporaceae | 7 | 20 | 6 | 50 | 100 | 67 | 30 | 29 | 88 | 11 | | 38 |
| **1145** | ***Pleosporales*** sp. | Ascomycota | Dothideomycetes | Pleosporales | | | | | | | 17 | | | | 89 | 100 | 88 |

| | | | | | | | | | | | | | | | | | |
|---|---|---|---|---|---|---|---|---|---|---|---|---|---|---|---|---|---|
| **672** | ***Dothideomycetes*** sp. | Ascomycota | Dothideomycetes | | | | 40 | 100 | | | | | | | | | |
| **1943** | ***Helotiales*** sp. | Ascomycota | Leotiomycetes | Helotiales | | 100 | 100 | 94 | | | 17 | 20 | 29 | 25 | 33 | 17 | 38 |
| **246** | ***Saccharomyces cerevisiae*** | Ascomycota | Saccharomycetes | Saccharomycetales | Saccharomycetaceae | 40 | 60 | 94 | | | | 20 | 29 | 25 | 22 | 33 | 38 |
| **971** | ***Clonostachys rosea*** | Ascomycota | Sordariomycetes | Hypocreales | Bionectriaceae | 93 | 60 | | 67 | 83 | 67 | 20 | 29 | | | | 25 |
| **1171** | ***Fusarium langsethiae*** | Ascomycota | Sordariomycetes | Hypocreales | Nectriaceae | 7 | 80 | 75 | | | | | 14 | | | | |
| **1167** | ***Gibberella fujikuroi*** | Ascomycota | Sordariomycetes | Hypocreales | Nectriaceae | 100 | 80 | 31 | 100 | 50 | 67 | 50 | 86 | 63 | 22 | 17 | 88 |
| **1113** | ***Gibberella tricincta*** | Ascomycota | Sordariomycetes | Hypocreales | Nectriaceae | 93 | 100 | 100 | 17 | 17 | | 50 | 57 | | 22 | | |
| **1394** | ***Ilyonectria macrodidyma*** | Ascomycota | Sordariomycetes | Hypocreales | Nectriaceae | 7 | | 13 | 83 | 50 | 83 | 60 | 29 | | | | 13 |
| **1814** | ***Nigrospora oryzae*** | Ascomycota | Sordariomycetes | Trichosphaeriales | Trichosphaeriaceae | | | | 50 | | 83 | | | 13 | | | |
| **892** | ***Xylariales*** **sp.** | Ascomycota | Sordariomycetes | Xylariales | | 100 | 60 | | 33 | 33 | 33 | 10 | | 25 | | | |
| **951** | ***Sordariomycetes*** sp. | Ascomycota | Sordariomycetes | | | | | | | | | 90 | 100 | 100 | 44 | 67 | 50 |
| **1195** | ***Pezizomycotina*** sp. 1 | Ascomycota | | | | 100 | 60 | 6 | 33 | 33 | 17 | | | 25 | | 17 | 50 |
| **1650** | ***Pezizomycotina*** sp. 2 | Ascomycota | | | | 100 | 80 | 100 | | | | 20 | 29 | 38 | 11 | | 13 |
| **489** | ***Bolbitius coprophilus*** | Basidiomycota | Agaricomycetes | Agaricales | Bolbitiaceae | | 20 | 94 | | | | | | | | | |
| **460** | ***Pholiotina brunnea*** | Basidiomycota | Agaricomycetes | Agaricales | Bolbitiaceae | 7 | 40 | 94 | | | | | | | | | |
| **1062** | ***Inocybe leucoloma*** | Basidiomycota | Agaricomycetes | Agaricales | Inocybaceae | 73 | 20 | 81 | | 33 | | 10 | 14 | | 22 | 33 | 50 |
| **276** | ***Inocybe malenconii*** | Basidiomycota | Agaricomycetes | Agaricales | Inocybaceae | 27 | 40 | 100 | | | | | | 13 | | 33 | 13 |
| **168** | ***Cylindrobasidium evolvens*** | Basidiomycota | Agaricomycetes | Agaricales | Physalacriaceae | | 20 | 100 | | | | | 14 | | | | |
| **1093** | ***Atheliaceae*** sp. | Basidiomycota | Agaricomycetes | Atheliales | Atheliaceae | | 40 | 81 | | | | | | | | | |
| **295** | ***Hypholoma sublateritium*** | Basidiomycota | Agaricomycetes | Atheliales | Strophariaceae | | 40 | 100 | | | | 10 | | | 11 | 50 | 13 |
| **941** | ***Sistotrema*** sp. | Basidiomycota | Agaricomycetes | Cantharellales | | 93 | 40 | 56 | | | | | 14 | 13 | | | |
| **290** | ***Sistotrema brinkmannii*** | Basidiomycota | Agaricomycetes | Cantharellales | | | 40 | 88 | | | | | | | | | |
| **500** | ***Corticium*** sp. | Basidiomycota | Agaricomycetes | Corticiales | Corticiaceae | | 40 | 100 | | | | | | | | | 25 |
| **594** | ***Vuilleminia comedens*** | Basidiomycota | Agaricomycetes | Corticiales | Vuilleminiaceae | | | | 17 | | 17 | 10 | 29 | 13 | 89 | 100 | 88 |
| **733** | ***Bjerkandera adusta*** | Basidiomycota | Agaricomycetes | Polyporales | Meruliaceae | 93 | 100 | 100 | 67 | 17 | 100 | 100 | 100 | 100 | 100 | 100 | 100 |
| **451** | ***Daedaleopsis confragosa*** | Basidiomycota | Agaricomycetes | Polyporales | Polyporaceae | 7 | 80 | 100 | | | | 30 | 86 | 100 | 67 | 67 | 88 |
| **479** | ***Lentinus tuber-regium*** | Basidiomycota | Agaricomycetes | Polyporales | Polyporaceae | 60 | 60 | 81 | | | | | | | | | |
| **804** | ***Trametes versicolor*** | Basidiomycota | Agaricomycetes | Polyporales | Polyporaceae | 13 | 40 | 100 | | | 17 | | | | | | 25 |
| **473** | ***Polyporales*** sp. | Basidiomycota | Agaricomycetes | Polyporales | | | 40 | 100 | | | | | | | 22 | | |

| | | | | | | | | | | | | | | | | | |
|---|---|---|---|---|---|---|---|---|---|---|---|---|---|---|---|---|---|
| **517** | ***Peniophora incarnata*** | Basidiomycota | Agaricomycetes | Russulales | Peniophoraceae | | 40 | 100 | 17 | | 17 | | | | | | |
| **985** | ***Kurtzmanomyces* sp.** | Basidiomycota | Agaricostilbomycetes | Agaricostilbales | Chionosphaeraceae | 87 | 20 | 75 | | | 17 | | | | | | 13 |
| **265** | ***Cystofilobasidium capitatum*** | Basidiomycota | Tremellomycetes | Cystofilobasidiales | Cystofilobasidiaceae | 20 | 60 | 88 | | | | | 14 | 13 | | | 13 |
| **937** | ***Cutaneotrichosporon cyanovorans*** | Basidiomycota | Tremellomycetes | Trichosporonales | Trichosporonaceae | 100 | 60 | 13 | | 17 | 17 | 10 | | | | | 13 |
| **165** | ***Malassezia globosa*** | Basidiomycota | Ustilaginomycotina Incertae sedis | Malasseziales | Malasseziaceae | 33 | 40 | 94 | | | 33 | 90 | 100 | 88 | 22 | 50 | 63 |
| **178** | ***Malassezia restricta*** | Basidiomycota | Ustilaginomycotina Incertae sedis | Malasseziales | Malasseziaceae | 53 | 80 | 94 | 17 | | 50 | 70 | 100 | 100 | 33 | 50 | 88 |
| **289** | ***Wallemia muriae*** | Basidiomycota | Wallemiomycetes | Wallemiales | Wallemiaceae | | 40 | 100 | | | | 10 | | 38 | | | 13 |
| **209** | ***Bifiguratus adelaidae*** | Mucoromycota | Endogonomycetes | Endogonales | | | | | | | | 40 | 43 | 38 | 56 | 67 | 88 |

**Table S2.** PERMANOVA results assessing the effects of species, organ, and their interaction on the Bray-Curtis and Jaccard dissimilarity matrixes of the OrM fungi dataset. The analysis was conducted using 9999 permutations to determine the significance of each term, revealing that species, organ, and their interaction significantly influence the community composition, as indicated by the F-values and $R^2$ values.

| | Term | Df | Sum Sq | Mean Sq | F value | $R^2$ | Significance |
|---|---|---|---|---|---|---|---|
| **Bray-Curtis** | Species | 3 | 8.9200 | 2.9733 | 11.5164 | 0.3368 | *** |
| | Organ | 2 | 1.3408 | 0.6704 | 2.5966 | 0.0506 | *** |
| | Species:Organ | 6 | 3.0592 | 0.5099 | 1.9749 | 0.1155 | *** |
| | Residuals | 51 | 13.1673 | 0.2582 | NA | 0.4971 | |
| | Total | 62 | 26.4873 | NA | NA | 1.0000 | |
| **Jaccard** | Species | 3 | 9.5372 | 3.1791 | 27.3354 | 0.5559 | 0.0001 *** |
| | Organ | 2 | 0.5865 | 0.2933 | 2.5217 | 0.0342 | 0.0053 ** |
| | Species:Organ | 6 | 1.1001 | 0.1834 | 1.5766 | 0.0641 | 0.0259 * |
| | Residuals | 51 | 5.9312 | 0.1163 | NA | 0.3457 | |
| | Total | 62 | 17.1551 | NA | NA | 1.0000 | |

**Table S3.** Summary of Constrained Analysis of Principal Coordinates (CAP) results comparing the effects of species identity, organ type (vegetative or reproductive), and their interaction on OrM fungal community composition. The table shows the F-statistic, $R^2$ (proportion of variance explained), p-values, and significance levels for each factor and their interaction. Analyses were performed using both Bray-Curtis and Jaccard distance metrics. Significance levels are indicated as: *** ($p < 0.001$), ** ($p < 0.01$), * ($p < 0.05$), ns (not significant). The results show that species identity and the interaction between species and organ type have a highly significant effect on OrM fungal community composition ($p < 0.001$) with both distance metrics.

| Analysis | F_statistic | $R^2$ | P_value | Significance |
|---|---|---|---|---|
| Species (Bray-Curtis) | 11.396 | 0.279 | 0.0001 | *** |
| Species (Jaccard) | 8.3563 | 0.221 | 0.0001 | *** |
| Organ Type (Bray-Curtis) | 1.6397 | 0.0266 | 0.0879 | ns |
| Organ Type (Jaccard) | 1.7335 | 0.0281 | 0.0509 | ns |
| Species × Organ Type (Bray-Curtis) | 5.4148 | 0.326 | 0.0001 | *** |
| Species × Organ Type (Jaccard) | 4.3352 | 0.279 | 0.0001 | *** |

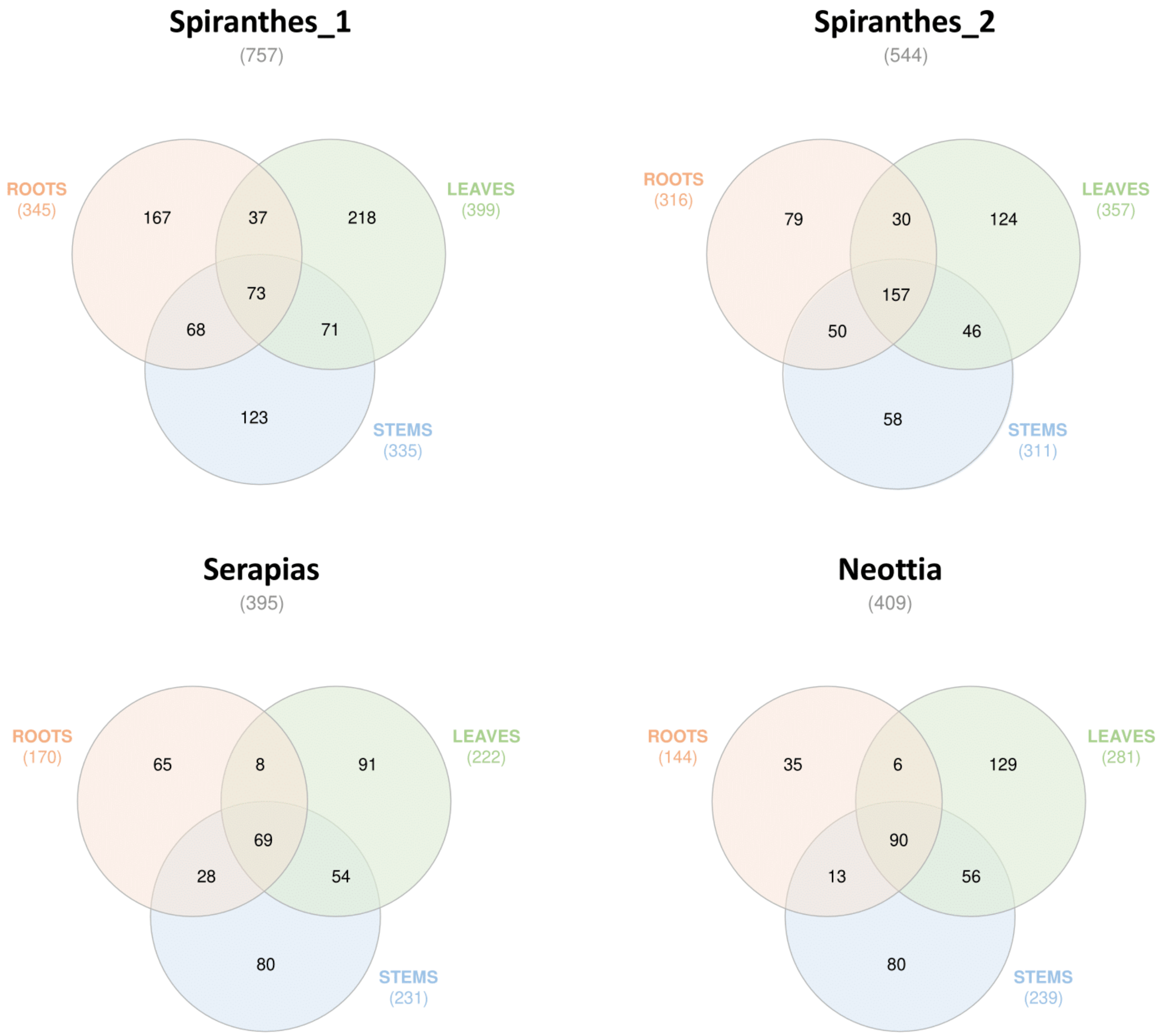


**Supplementary figure S1**. Venn diagrams showing the number of shared and organ-specific fungal OTUs in the roots, stems and leaves of the same plant for each of the three orchid species: *Spiranthes spiralis*, *Serapias vomeracea* and *Neottia ovata*. For *S. spiralis*, plants were sampled in two consecutive years and the data were kept separate.

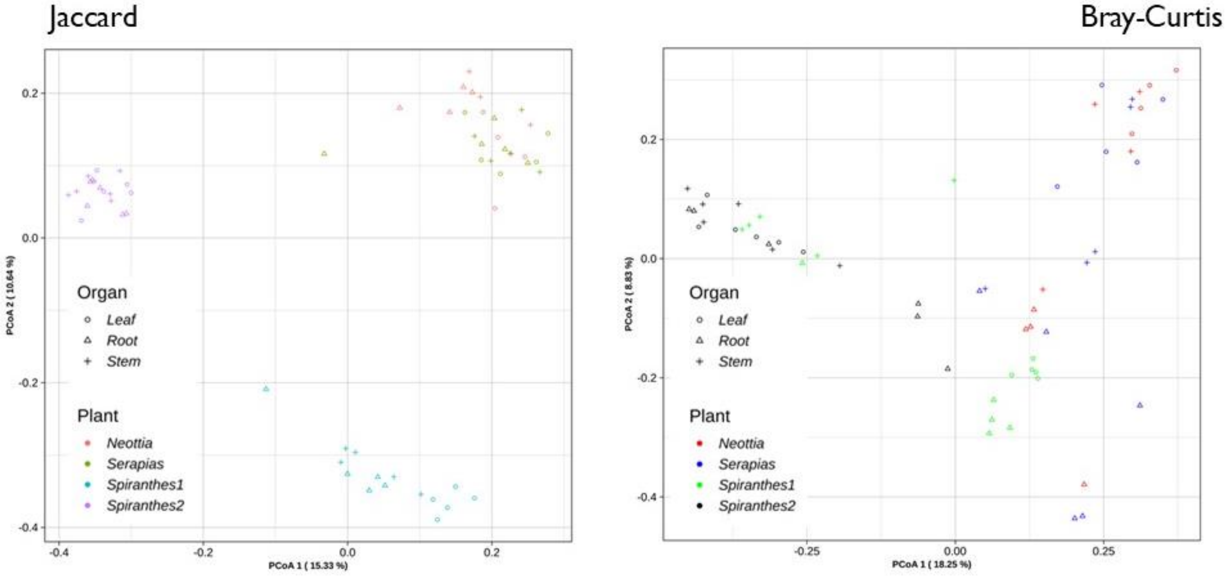


**Supplementary figure S2.** Principal Component Analysis (PCA) of fungal communities associated with different orchid organs based on Jaccard (left) and Bray-Curtis (right) dissimilarity matrices. The analysis includes fungal OTUs from three orchid taxa (*Neottia ovata*, *Serapias vomeracea*, and two *Spiranthes spiralis* sample groups) and three plant organs (leaf, root, and stem). Points are shaped according to the organ type (circles = leaf, triangles = root, crosses = stem) and colored by orchid taxa. The percentage of variance explained by each principal component is shown on the axes. PCoA1 explains 15.33% and 18.25% of the total variance for Jaccard and Bray-Curtis analyses respectively.

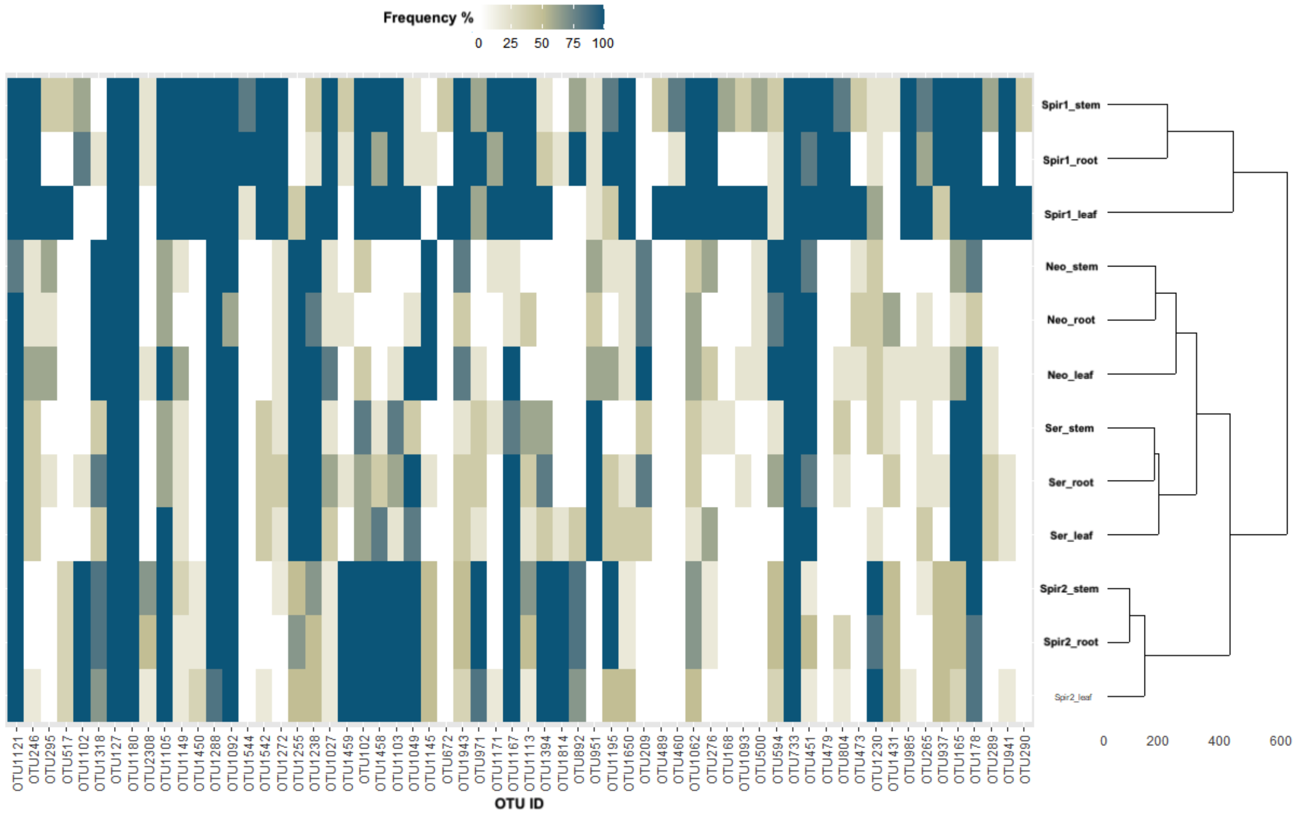


**Supplementary figure S3**. Heatmap and hierarchical clustering of the dominant non-OrM fungal OTUs across plant species and organs. The heatmap displays OTU frequency percentages (0-100%) in different plant organs (stem, root, leaf) of the target orchids: *Spiranthes spiralis_1* (Spir1), *Spiranthes spiralis_2* (Spir2), *Neottia ovata* (Neo), *Serapias vomeracea* (Ser). Hierarchical clustering (dendrogram) shows sample relationships. Color intensity from white to dark blue represents increasing OTU frequency.

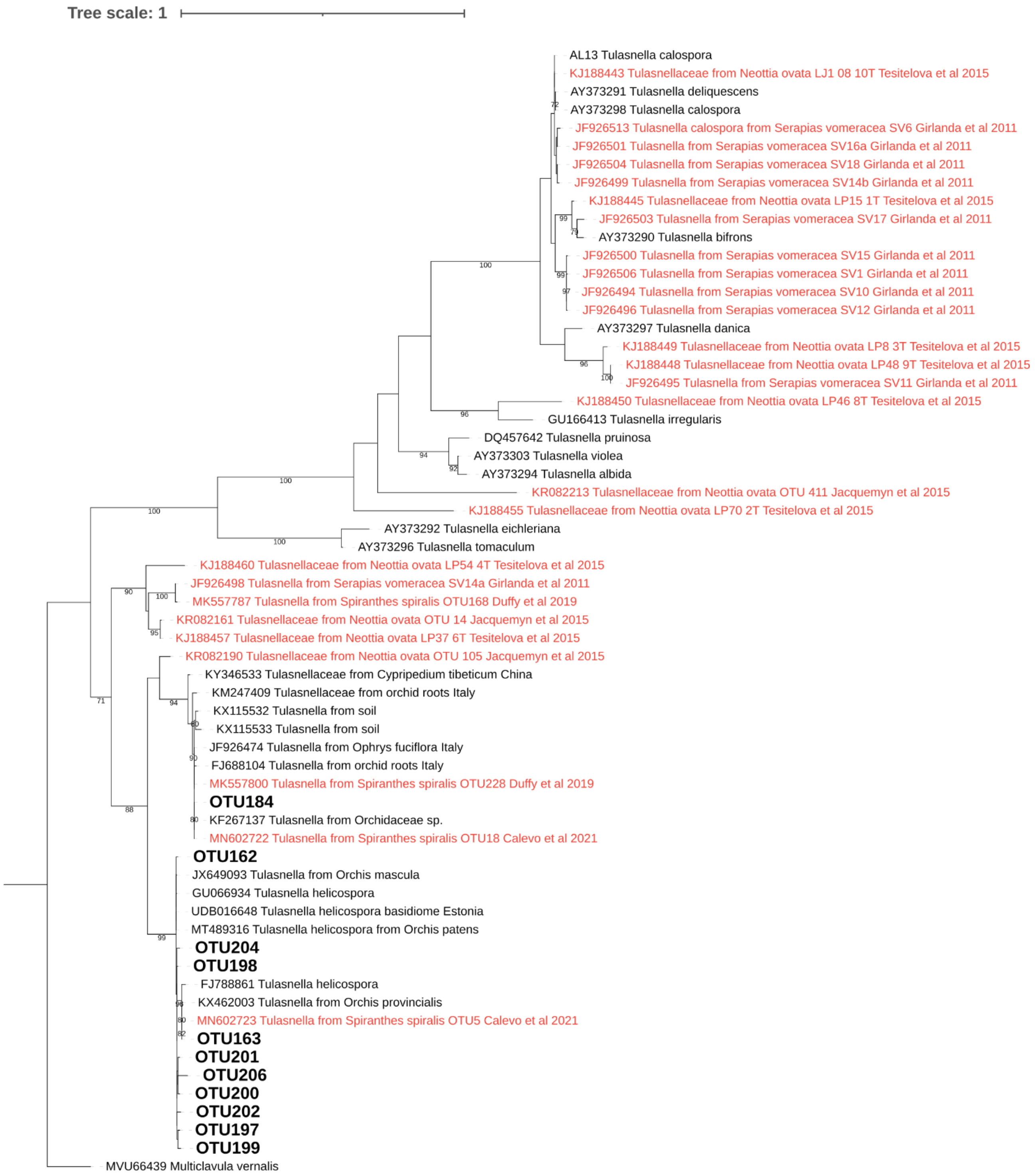


**Supplementary figure S4**. Maximum likelihood tree obtained from the ITS2 sequence alignment of tulasnelloid fungi. *Multicalvula vernalis* was used as outgroup. Bootstrap support values above 70% (1000 maximum likelihood replicates) are reported. OTUs found in this study are indicated in bald, sequences retrieved in previous studies from the same orchid species are in red.

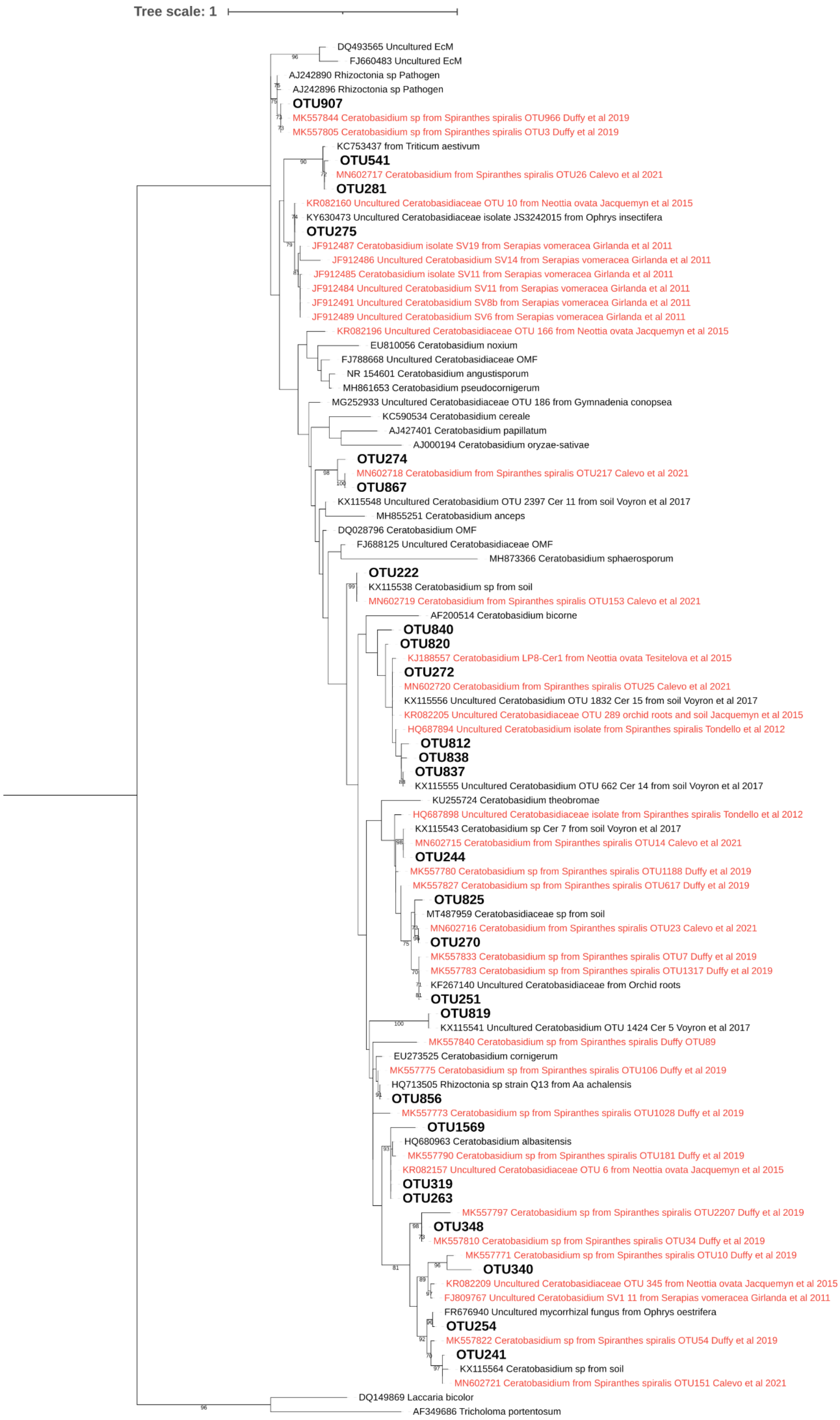

Tree scale: 1
DQ493565 Uncultured EcM
FJ660483 Uncultured EcM
AJ242890 Rhizoctonia sp Pathogen
AJ242896 Rhizoctonia sp Pathogen
OTU907
MK557844 Ceratobasidium sp from Spiranthes spiralis OTU966 Duffy et al 2019
MK557805 Ceratobasidium sp from Spiranthes spiralis OTU3 Duffy et al 2019
KC753437 from Triticum aestivum
OTU541
MN602717 Ceratobasidium from Spiranthes spiralis OTU26 Calevo et al 2021
OTU281
KR082160 Uncultured Ceratobasidiaceae OTU 10 from Neottia ovata Jacquemyn et al 2015
KY630473 Uncultured Ceratobasidiaceae isolate JS3242015 from Ophrys insectifera
OTU275
JF912487 Ceratobasidium isolate SV19 from Serapias vomeracea Girlanda et al 2011
JF912486 Uncultured Ceratobasidium SV14 from Serapias vomeracea Girlanda et al 2011
JF912485 Ceratobasidium isolate SV11 from Serapias vomeracea Girlanda et al 2011
JF912484 Uncultured Ceratobasidium SV11 from Serapias vomeracea Girlanda et al 2011
JF912491 Uncultured Ceratobasidium SV8b from Serapias vomeracea Girlanda et al 2011
JF912489 Uncultured Ceratobasidium SV6 from Serapias vomeracea Girlanda et al 2011
KR082196 Uncultured Ceratobasidiaceae OTU 166 from Neottia ovata Jacquemyn et al 2015
EU810056 Ceratobasidium noxium
FJ788668 Uncultured Ceratobasidiaceae OMF
NR 154601 Ceratobasidium angustisporum
MH861653 Ceratobasidium pseudocornigerum
MG252933 Uncultured Ceratobasidiaceae OTU 186 from Gymnadenia conopsea
KC590534 Ceratobasidium cereale
AJ427401 Ceratobasidium papillatum
AJ000194 Ceratobasidium oryzae-sativae
OTU274
MN602718 Ceratobasidium from Spiranthes spiralis OTU217 Calevo et al 2021
OTU867
KX115548 Uncultured Ceratobasidium OTU 2397 Cer 11 from soil Voyron et al 2017
MH855251 Ceratobasidium anceps
DQ028796 Ceratobasidium OMF
FJ688125 Uncultured Ceratobasidiaceae OMF
MH873366 Ceratobasidium sphaerosporum
OTU222
KX115538 Ceratobasidium sp from soil
MN602719 Ceratobasidium from Spiranthes spiralis OTU153 Calevo et al 2021
AF200514 Ceratobasidium bicorne
OTU840
OTU820
KJ188557 Ceratobasidium LP8-Cer1 from Neottia ovata Tesitelova et al 2015
OTU272
MN602720 Ceratobasidium from Spiranthes spiralis OTU25 Calevo et al 2021
KX115556 Uncultured Ceratobasidium OTU 1832 Cer 15 from soil Voyron et al 2017
KR082205 Uncultured Ceratobasidiaceae OTU 289 orchid roots and soil Jacquemyn et al 2015
HQ687894 Uncultured Ceratobasidium isolate from Spiranthes spiralis Tondello et al 2012
OTU812
OTU838
OTU837
KX115555 Uncultured Ceratobasidium OTU 662 Cer 14 from soil Voyron et al 2017
KU255724 Ceratobasidium theobromae
HQ687898 Uncultured Ceratobasidiaceae isolate from Spiranthes spiralis Tondello et al 2012
KX115543 Ceratobasidium sp Cer 7 from soil Voyron et al 2017
MN602715 Ceratobasidium from Spiranthes spiralis OTU14 Calevo et al 2021
OTU244
MK557780 Ceratobasidium sp from Spiranthes spiralis OTU1188 Duffy et al 2019
MK557827 Ceratobasidium sp from Spiranthes spiralis OTU617 Duffy et al 2019
OTU825
MT487959 Ceratobasidiaceae sp from soil
MN602716 Ceratobasidium from Spiranthes spiralis OTU23 Calevo et al 2021
OTU270
MK557833 Ceratobasidium sp from Spiranthes spiralis OTU7 Duffy et al 2019
MK557783 Ceratobasidium sp from Spiranthes spiralis OTU1317 Duffy et al 2019
KF267140 Uncultured Ceratobasidiaceae from Orchid roots
OTU251
OTU819
KX115541 Uncultured Ceratobasidium OTU 1424 Cer 5 Voyron et al 2017
MK557840 Ceratobasidium sp from Spiranthes spiralis Duffy OTU89
EU273525 Ceratobasidium cornigerum
MK557775 Ceratobasidium sp from Spiranthes spiralis OTU106 Duffy et al 2019
HQ713505 Rhizoctonia sp strain Q13 from Aa achalensis
OTU856
MK557773 Ceratobasidium sp from Spiranthes spiralis OTU1028 Duffy et al 2019
OTU1569
HQ680963 Ceratobasidium albasitensis
MK557790 Ceratobasidium sp from Spiranthes spiralis OTU181 Duffy et al 2019
KR082157 Uncultured Ceratobasidiaceae OTU 6 from Neottia ovata Jacquemyn et al 2015
OTU319
OTU263
MK557797 Ceratobasidium sp from Spiranthes spiralis OTU2207 Duffy et al 2019
OTU348
MK557810 Ceratobasidium sp from Spiranthes spiralis OTU34 Duffy et al 2019
MK557771 Ceratobasidium sp from Spiranthes spiralis OTU10 Duffy et al 2019
OTU340
KR082209 Uncultured Ceratobasidiaceae OTU 345 from Neottia ovata Jacquemyn et al 2015
FJ809767 Uncultured Ceratobasidium SV1 11 from Serapias vomeracea Girlanda et al 2011
FR676940 Uncultured mycorrhizal fungus from Ophrys oestrifera
OTU254
MK557822 Ceratobasidium sp from Spiranthes spiralis OTU54 Duffy et al 2019
OTU241
KX115564 Ceratobasidium sp from soil
MN602721 Ceratobasidium from Spiranthes spiralis OTU151 Calevo et al 2021
DQ149869 Laccaria bicolor
AF349686 Tricholoma portentosum

**Supplementary figure S5**. Maximum likelihood tree obtained from the ITS2 sequence alignment of ceratobasidioid fungi. *Tricholoma portentosum* and *Laccaria bicolor* were used as outgroup taxa. Bootstrap support values above 70% (1000 maximum likelihood replicates) are reported. OTUs found in this study are indicated in bald, sequences retrieved in previous studies from the same orchid species are in red.

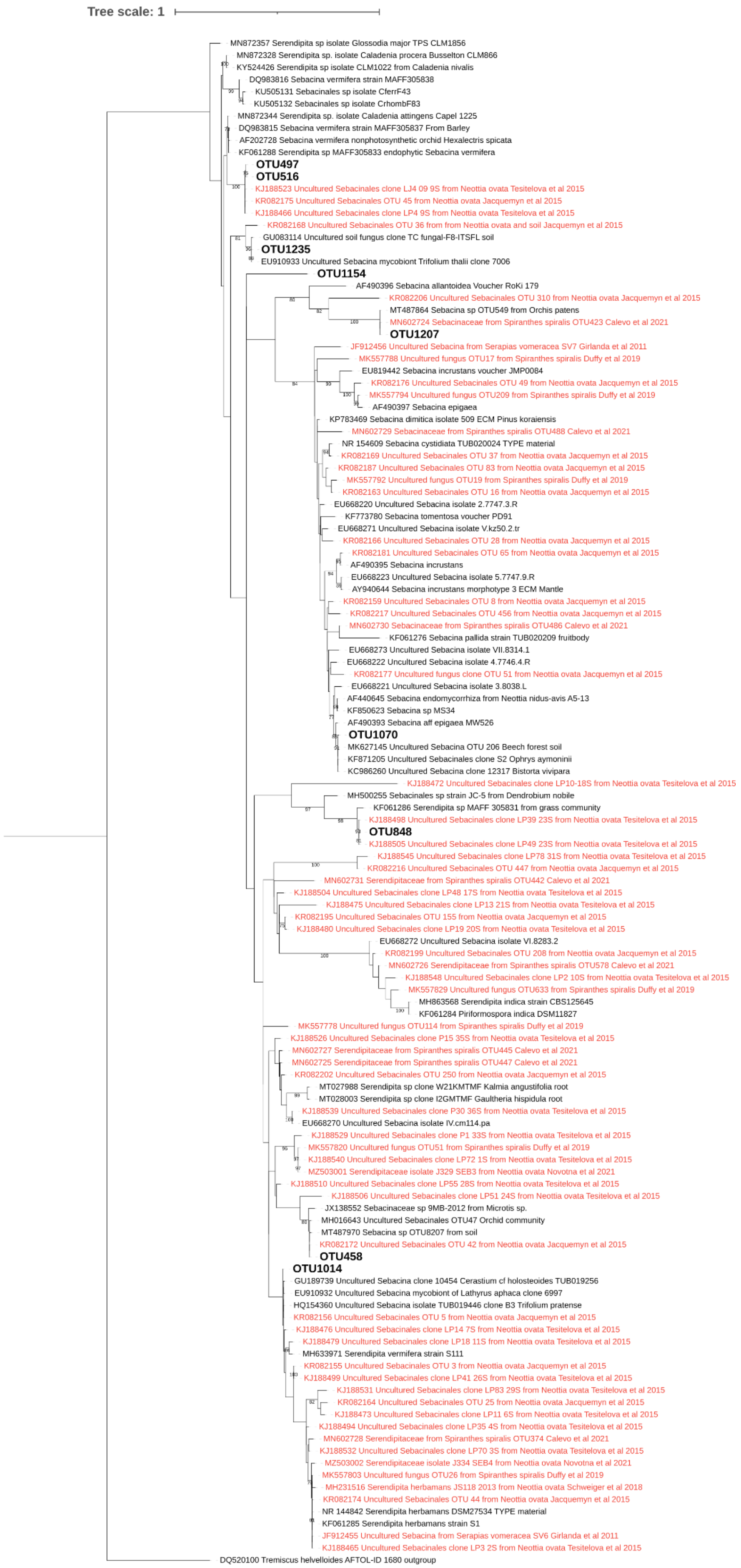
Tree scale: 1
MN872357 Serendipita sp isolate Glossodia major TPS CLM1856
MN872328 Serendipita sp. isolate Caladenia procera Busselton CLM866
KY524426 Serendipita sp isolate CLM1022 from Caladenia nivalis
DQ983816 Sebacina vermifera strain MAFF305838
KU505131 Sebacinales sp isolate CferrF43
KU505132 Sebacinales sp isolate CrhombF83
MN872344 Serendipita sp. isolate Caladenia attingens Capel 1225
DQ983815 Sebacina vermifera strain MAFF305837 From Barley
AF202728 Sebacina vermifera nonphotosynthetic orchid Hexalectris spicata
KF061288 Serendipita sp MAFF305833 endophytic Sebacina vermifera
OTU497
OTU516
KJ188523 Uncultured Sebacinales clone LJ4 09 9S from Neottia ovata Tesitelova et al 2015
KR082175 Uncultured Sebacinales OTU 45 from Neottia ovata Jacquemyn et al 2015
KJ188466 Uncultured Sebacinales clone LP4 9S from Neottia ovata Tesitelova et al 2015
KR082168 Uncultured Sebacinales OTU 36 from from Neottia ovata and soil Jacquemyn et al 2015
GU083114 Uncultured soil fungus clone TC fungal-F8-ITSFL soil
OTU1235
EU910933 Uncultured Sebacina mycobiont Trifolium thalii clone 7006
OTU1154
AF490396 Sebacina allantoidea Voucher RoKi 179
KR082206 Uncultured Sebacinales OTU 310 from Neottia ovata Jacquemyn et al 2015
MT487864 Sebacina sp OTU549 from Orchis patens
MN602724 Sebacinaceae from Spiranthes spiralis OTU423 Calevo et al 2021
OTU1207
JF912456 Uncultured Sebacina from Serapias vomeracea SV7 Girlanda et al 2011
MK557788 Uncultured fungus OTU17 from Spiranthes spiralis Duffy et al 2019
EU819442 Sebacina incrustans voucher JMP0084
KR082176 Uncultured Sebacinales OTU 49 from Neottia ovata Jacquemyn et al 2015
MK557794 Uncultured fungus OTU209 from Spiranthes spiralis Duffy et al 2019
AF490397 Sebacina epigaea
KP783469 Sebacina dimitica isolate 509 ECM Pinus koraiensis
MN602729 Sebacinaceae from Spiranthes spiralis OTU488 Calevo et al 2021
NR 154609 Sebacina cystidiata TUB020024 TYPE material
KR082169 Uncultured Sebacinales OTU 37 from Neottia ovata Jacquemyn et al 2015
KR082187 Uncultured Sebacinales OTU 83 from Neottia ovata Jacquemyn et al 2015
MK557792 Uncultured fungus OTU19 from Spiranthes spiralis Duffy et al 2019
KR082163 Uncultured Sebacinales OTU 16 from Neottia ovata Jacquemyn et al 2015
EU668220 Uncultured Sebacina isolate 2.7747.3.R
KF773780 Sebacina tomentosa voucher PD91
EU668271 Uncultured Sebacina isolate V.kz50.2.tr
KR082166 Uncultured Sebacinales OTU 28 from Neottia ovata Jacquemyn et al 2015
KR082181 Uncultured Sebacinales OTU 65 from Neottia ovata Jacquemyn et al 2015
AF490395 Sebacina incrustans
EU668223 Uncultured Sebacina isolate 5.7747.9.R
AY940644 Sebacina incrustans morphotype 3 ECM Mantle
KR082159 Uncultured Sebacinales OTU 8 from Neottia ovata Jacquemyn et al 2015
KR082217 Uncultured Sebacinales OTU 456 from Neottia ovata Jacquemyn et al 2015
MN602730 Sebacinaceae from Spiranthes spiralis OTU486 Calevo et al 2021
KF061276 Sebacina pallida strain TUB020209 fruitbody
EU668273 Uncultured Sebacina isolate VII.8314.1
EU668222 Uncultured Sebacina isolate 4.7746.4.R
KR082177 Uncultured fungus clone OTU 51 from Neottia ovata Jacquemyn et al 2015
EU668221 Uncultured Sebacina isolate 3.8038.L
AF440645 Sebacina endomycorrhiza from Neottia nidus-avis A5-13
KF850623 Sebacina sp MS34
AF490393 Sebacina aff epigaea MW526
OTU1070
MK627145 Uncultured Sebacina OTU 206 Beech forest soil
KF871205 Uncultured Sebacinales clone S2 Ophrys aymoninii
KC986260 Uncultured Sebacina clone 12317 Bistorta vivipara
KJ188472 Uncultured Sebacinales clone LP10-18S from Neottia ovata Tesitelova et al 2015
MH500255 Sebacinales sp strain JC-5 from Dendrobium nobile
KF061286 Serendipita sp MAFF 305831 from grass community
KJ188498 Uncultured Sebacinales clone LP39 23S from Neottia ovata Tesitelova et al 2015
OTU848
KJ188505 Uncultured Sebacinales clone LP49 23S from Neottia ovata Tesitelova et al 2015
KJ188545 Uncultured Sebacinales clone LP78 31S from Neottia ovata Tesitelova et al 2015
KR082216 Uncultured Sebacinales OTU 447 from Neottia ovata Jacquemyn et al 2015
MN602731 Serendipitaceae from Spiranthes spiralis OTU442 Calevo et al 2021
KJ188504 Uncultured Sebacinales clone LP48 17S from Neottia ovata Tesitelova et al 2015
KJ188475 Uncultured Sebacinales clone LP13 21S from Neottia ovata Tesitelova et al 2015
KR082195 Uncultured Sebacinales OTU 155 from Neottia ovata Jacquemyn et al 2015
KJ188480 Uncultured Sebacinales clone LP19 20S from Neottia ovata Tesitelova et al 2015
EU668272 Uncultured Sebacina isolate VI.8283.2
KR082199 Uncultured Sebacinales OTU 208 from Neottia ovata Jacquemyn et al 2015
MN602726 Serendipitaceae from Spiranthes spiralis OTU578 Calevo et al 2021
KJ188548 Uncultured Sebacinales clone LP2 10S from Neottia ovata Tesitelova et al 2015
MK557829 Uncultured fungus OTU633 from Spiranthes spiralis Duffy et al 2019
MH863568 Serendipita indica strain CBS125645
KF061284 Piriformospora indica DSM11827
MK557778 Uncultured fungus OTU114 from Spiranthes spiralis Duffy et al 2019
KJ188526 Uncultured Sebacinales clone P15 35S from Neottia ovata Tesitelova et al 2015
MN602727 Serendipitaceae from Spiranthes spiralis OTU445 Calevo et al 2021
MN602725 Serendipitaceae from Spiranthes spiralis OTU447 Calevo et al 2021
KR082202 Uncultured Sebacinales OTU 250 from Neottia ovata Jacquemyn et al 2015
MT027988 Serendipita sp clone W21KMTMF Kalmia angustifolia root
MT028003 Serendipita sp clone I2GMTMF Gaultheria hispidula root
KJ188539 Uncultured Sebacinales clone P30 36S from Neottia ovata Tesitelova et al 2015
EU668270 Uncultured Sebacina isolate IV.cm114.pa
KJ188529 Uncultured Sebacinales clone P1 33S from Neottia ovata Tesitelova et al 2015
MK557820 Uncultured fungus OTU51 from Spiranthes spiralis Duffy et al 2019
KJ188540 Uncultured Sebacinales clone LP72 1S from Neottia ovata Tesitelova et al 2015
MZ503001 Serendipitaceae isolate J329 SEB3 from Neottia ovata Novotna et al 2021
KJ188510 Uncultured Sebacinales clone LP55 28S from Neottia ovata Tesitelova et al 2015
KJ188506 Uncultured Sebacinales clone LP51 24S from Neottia ovata Tesitelova et al 2015
JX138552 Sebacinaceae sp 9MB-2012 from Microtis sp.
MH016643 Uncultured Sebacinales OTU47 Orchid community
MT487970 Sebacina sp OTU8207 from soil
KR082172 Uncultured Sebacinales OTU 42 from Neottia ovata Jacquemyn et al 2015
OTU458
OTU1014
GU189739 Uncultured Sebacina clone 10454 Cerastium cf holosteoides TUB019256
EU910932 Uncultured Sebacina mycobiont of Lathyrus aphaca clone 6997
HQ154360 Uncultured Sebacina isolate TUB019446 clone B3 Trifolium pratense
KR082156 Uncultured Sebacinales OTU 5 from Neottia ovata Jacquemyn et al 2015
KJ188476 Uncultured Sebacinales clone LP14 7S from Neottia ovata Tesitelova et al 2015
KJ188479 Uncultured Sebacinales clone LP18 11S from Neottia ovata Tesitelova et al 2015
MH633971 Serendipita vermifera strain S111
KR082155 Uncultured Sebacinales OTU 3 from Neottia ovata Jacquemyn et al 2015
KJ188499 Uncultured Sebacinales clone LP41 26S from Neottia ovata Tesitelova et al 2015
KJ188531 Uncultured Sebacinales clone LP83 29S from Neottia ovata Tesitelova et al 2015
KR082164 Uncultured Sebacinales OTU 25 from Neottia ovata Jacquemyn et al 2015
KJ188473 Uncultured Sebacinales clone LP11 6S from Neottia ovata Tesitelova et al 2015
KJ188494 Uncultured Sebacinales clone LP35 4S from Neottia ovata Tesitelova et al 2015
MN602728 Serendipitaceae from Spiranthes spiralis OTU374 Calevo et al 2021
KJ188532 Uncultured Sebacinales clone LP70 3S from Neottia ovata Tesitelova et al 2015
MZ503002 Serendipitaceae isolate J334 SEB4 from Neottia ovata Novotna et al 2021
MK557803 Uncultured fungus OTU26 from Spiranthes spiralis Duffy et al 2019
MH231516 Serendipita herbamans JS118 2013 from Neottia ovata Schweiger et al 2018
KR082174 Uncultured Sebacinales OTU 44 from Neottia ovata Jacquemyn et al 2015
NR 144842 Serendipita herbamans DSM27534 TYPE material
KF061285 Serendipita herbamans strain S1
JF912455 Uncultured Sebacina from Serapias vomeracea SV6 Girlanda et al 2011
KJ188465 Uncultured Sebacinales clone LP3 2S from Neottia ovata Tesitelova et al 2015
DQ520100 Tremiscus helvelloides AFTOL-ID 1680 outgroup

**Supplementary figure S6**. Maximum likelihood tree obtained from the ITS2 sequence alignment of fungi assigned to Sebacinales. *Tremiscus helvelloides* was used as an outgroup taxon. Bootstrap support values above 70% (1000 maximum likelihood replicates) are reported. OTUs found in this study are indicated in bald, sequences retrieved in previous studies from the same orchid species are in red.

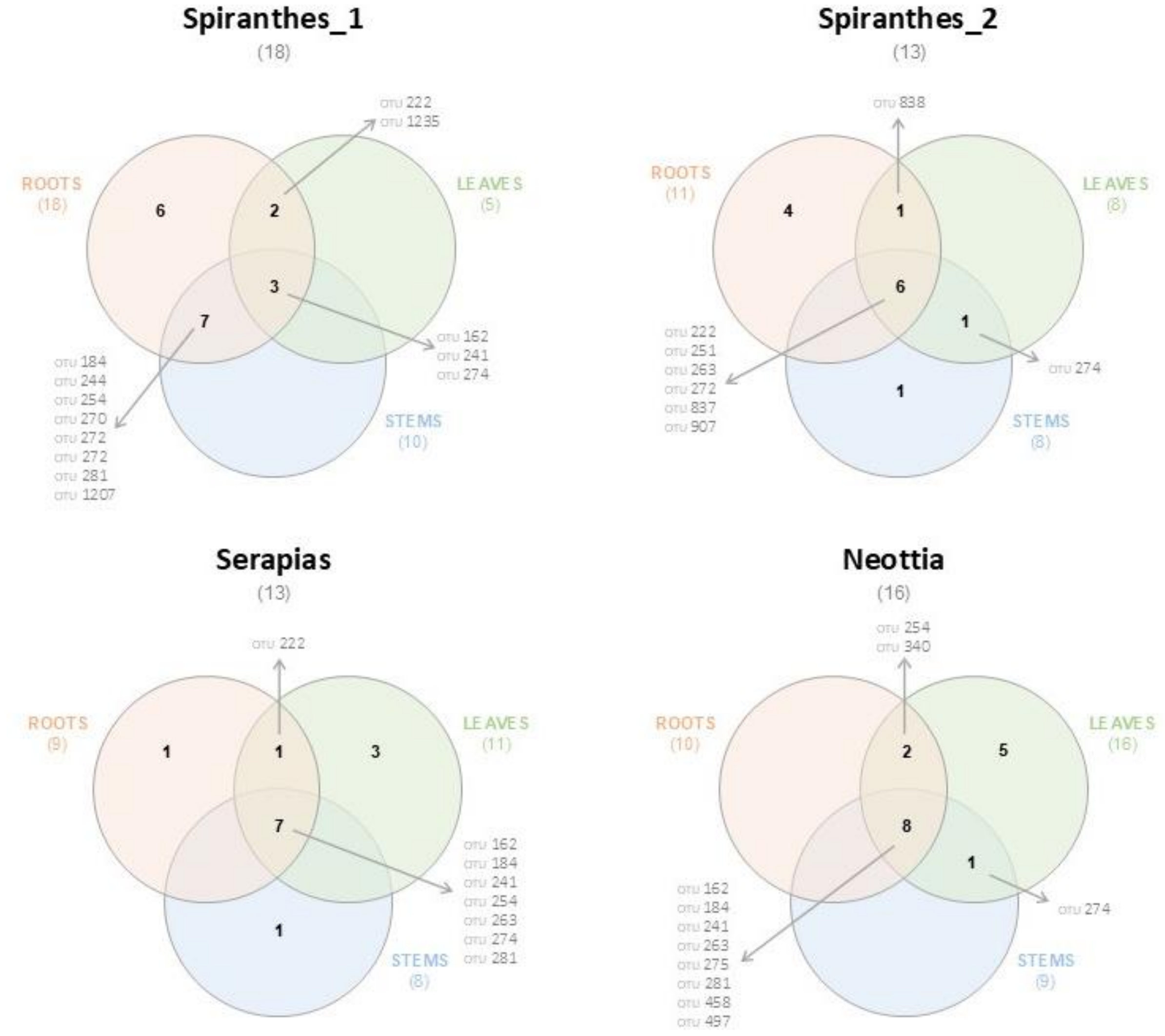


**Supplementary figure S7**. Venn diagrams showing the number of tulasnelloid, ceratobasidioid and sebacinoid s.l. OTUs in the roots, stems and leaves of the same plant for each of the three orchid species: *Spiranthes spiralis*, *Serapias vomeracea* and *Neottia ovata*. The list of OTUs shared between different organs is provided. For *S. spiralis*, plants were sampled in two consecutive years and results were kept separate.

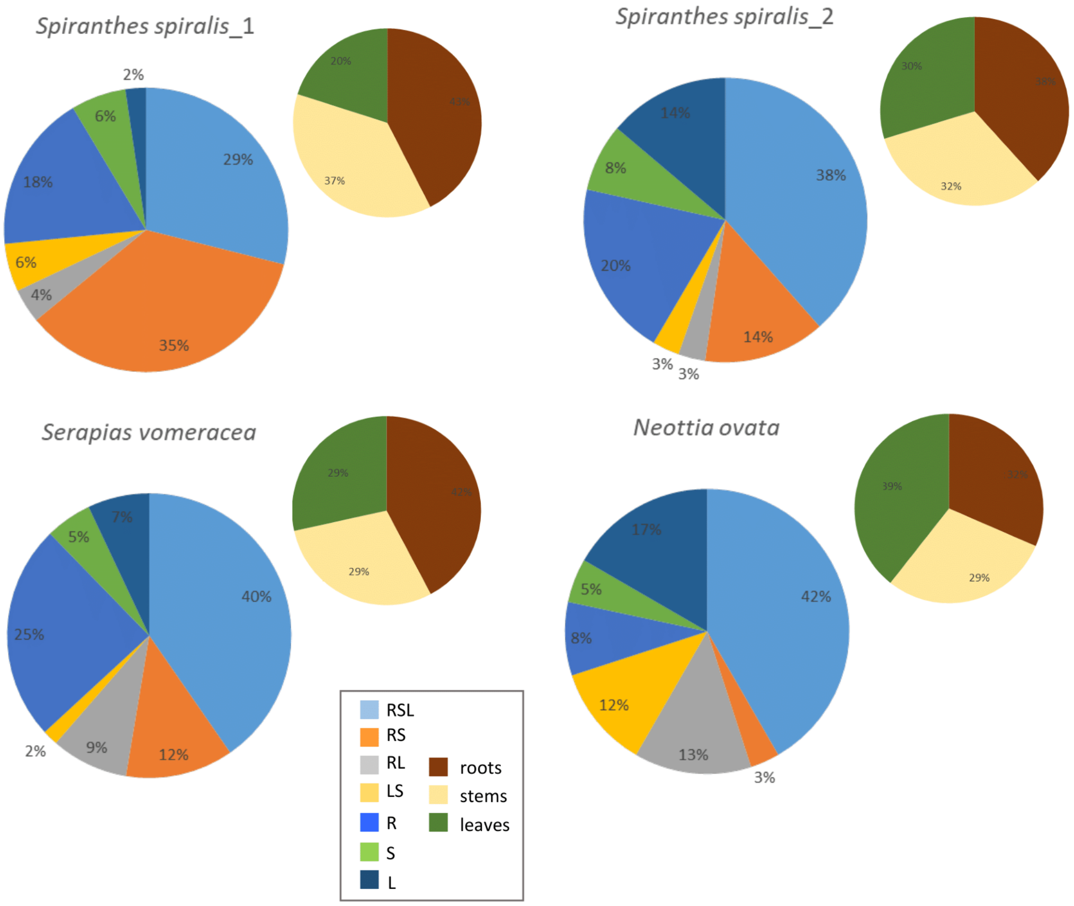


**Supplementary figure S8**. The larger pie charts depict the share of occurrences of the 33 rhizoctonia OTUs in either all (RSL), two (RS, roots&stem; RL, roots&leaves and LS, leaves&stem), or a single vegetative organ (R, roots; S, stem; L, leaves) in the same plant for each orchid species. The smaller charts summarize the share of total occurrences in roots, stems and leaves.

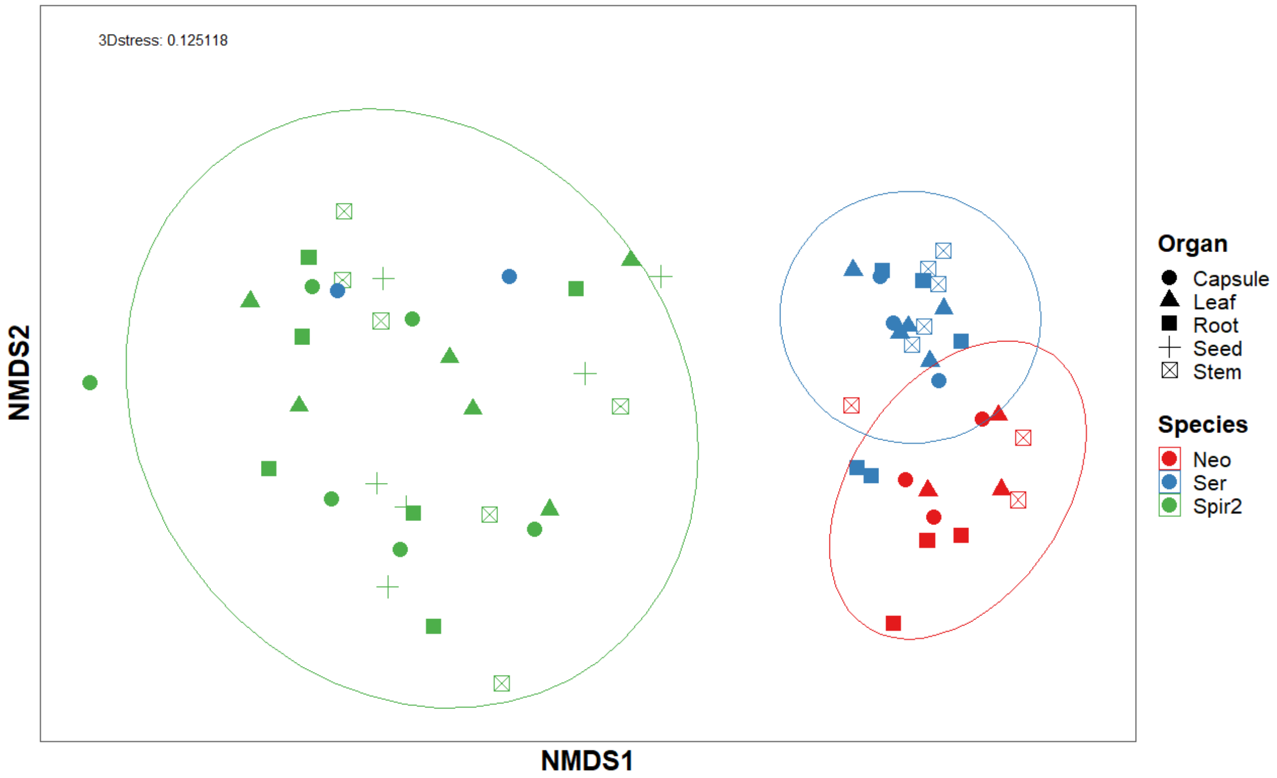


**Supplementary figure S9**. Non-metric multidimensional scaling (NMDS) plot illustrating the relationships among different plant organs and species based on their characteristics. The axes represent NMDS1 and NMDS2 dimensions. Data points are shaped according to organ type (capsule, leaf, root, seed, stem) and colored by species (*Neottia ovata*, *Serapias vomeracea*, *Spiranthes spiralis_2*). Ellipses indicate clustering patterns, with a stress value of 0.125118, suggesting a good fit for the ordination.
